# Systematic assessment of ISWI subunits reveals that NURF creates local accessibility for proper CTCF function

**DOI:** 10.1101/2023.07.25.550466

**Authors:** Mario Iurlaro, Francesca Masoni, Christiane Wirbelauer, Murat Iskar, Lukas Burger, Dirk Schübeler

## Abstract

Catalytic activity of the ISWI family of remodelers is critical for nucleosomal organization and DNA binding of transcription factors, including the insulator protein CTCF. To define which subcomplex mediates these diverse functions, we derived a panel of isogenic mouse stem cell lines each lacking one of six ISWI accessory subunits. Individual deletions of subunits of either CERF, RSF, ACF, WICH or NoRC subcomplexes only moderately affect the chromatin landscape, while removal of the NURF-specific subunit BPTF leads to drastic reduction in chromatin accessibility and SNF2H ATPase localization around CTCF sites. While this affects adjacent nucleosome occupancy, it only modestly impacts CTCF binding itself. In the absence of accessibility, the structural function of CTCF is nevertheless impaired resulting in lower occupancy of cohesin and cohesin release factor, and reduced physical insulation at these sites, highlighting the need of NURF-mediated remodeling for open chromatin and proper CTCF function.

These results separate local CTCF binding from insulator function in nuclear organization and reveal a specific role for NURF in mediating SNF2H localization and chromatin opening at bound CTCF sites. They designate local accessibility as critical for cohesin positioning and establishment of physical insulation.

## Introduction

The ability of chromatin remodeling enzymes to move, slide or evict nucleosomes is essential for all aspects of genome regulation^1^. This includes the ability of DNA binding factors to gain access to the genome, which is directly linked to presence and position of nucleosomes^2^. Indeed, sites bound by transcription factors (TFs) are characterized by high local accessibility, which is assumed to be required for their binding^3^.

How remodeling activity is targeted to specific genomic sites has been difficult to address due to the large number of potential complexes involved and the lack of a systematic assessment of their local contributions. Among these, ISWI represents one of the four mammalian remodeler families besides SWI/SNF, CHD and INO80^1^. ISWI family members consist of one of two exchangeable ATPases SNF2L (Smarca1) or SNF2H (Smarca5) that alternatively associate with one or more complex-specific accessory subunits. The main ISWI accessory subunits are RSF1, ACF1, WSTF, TIP5, CECR2 and BPTF, which together with one catalytic subunit create the RSF, ACF, WICH, NoRC, CERF or NURF complexes, respectively^4–11^. Additionally, some of these core complexes can associate with auxiliary components such as the histone-fold proteins CHRAC-15/17, which bind to ACF1-SNF2H to form the CHRAC complex^12^. Interestingly, the ability of SNF2H to sense its substrate in vitro differs within different complexes^13^. In vivo, ISWI complexes have distinct roles in transcription regulation including both activating and repressing mechanisms^4, 5^. The NURF, RSF, CERF and ACF complexes have been shown to impact transcription of genes transcribed by RNA Polymerase II, while NoRC and WICH complexes have been linked to RNA Polymerase I transcribed genes^4, 5^. Altogether, this suggests that ISWI complexes present distinct enzymatic abilities in vitro and are also associated with distinct functions in vivo. To explain complex-specific functions within the cell it has been proposed that accessory subunits could differentially regulate the ATPase activity in terms of localized recruitment or activation at specific chromatin regions^5^, however a systematic assessment of this model is presently missing.

Genetic deletion or targeted degradation of SNF2H in mammalian cells in culture causes genome-wide changes in nucleosome organization with increased nucleosomal repeat length and reduced binding of specific TFs, such as CTCF, which coincides with reduced chromatin accessibility^14, 15^. With the goal of assigning functions for individual ISWI subcomplexes we generated a set of isogenic murine stem cell lines with loss of function mutations of defined subunits. Detailed molecular phenotyping at the level of the transcriptome and the epigenetic landscape of accessibility and nucleosomal organization reveals that ACF1, RSF1, CECR2 and TIP5 only mildly contribute to nucleosomal positioning or transcription, possibly indicative of functional redundancy between the studied subcomplexes. However, we identify a specific role for the NURF subunit BPTF in generating accessible chromatin around CTCF bound sites. We show that, in the absence of chromatin opening via NURF, CTCF binding largely persists while its structural function as insulator is impaired. These findings link chromatin opening via NURF remodeling to CTCF insulator function.

## Results

### Generation of isogenic cell lines with individual ISWI deletions

Mammalian ISWI subcomplexes are considered mutually independent, with each consisting of a catalytic ATPase (either SNF2H or SNF2L) together with one or more non-catalytic accessory subunits. To characterize the protein composition of SNF2H-containing ISWI subcomplexes in mouse embryonic stem cells (mESCs), we performed co-immunoprecipitation in native conditions using the catalytic subunit SNF2H as bait (SNF2H co-IP) and quantified associated proteins by mass spectrometry using SNF2H depleted cells (Snf2hΔ) as a negative control^14^ (Methods). This revealed that SNF2H interacts with previously identified ISWI accessory subunits including CECR2 and BPTF that are more traditionally associated with SNF2L ^5, 16^ (Fig. 1a). In parallel, we also performed SNF2H co-IP followed by western blot detection for ISWI accessory subunits which, in addition to the previous approach, validates the previously reported interaction of SNF2H with WSTF and BAZ2A/TIP5 subunits in wild-type cells (WT lane in Extended Data Fig. 1a). While we detect BPTF by Mass Spectrometry upon SNF2H co-IP, this interaction proved to be more difficult to show via western blotting presumably due to the large size of the BPTF protein (predicted higher than 300 kDa) (wild-type lane in Fig. 1b and in Extended Data Fig. 1a). Taken together, this suggests that in mESCs, SNF2H not only interacts with its canonical partners (RSF1, TIP5, WSTF and ACF1) but also with CECR2 and BPTF (Fig. 1a, Extended Data Fig. 1a).

**Figure 1.**
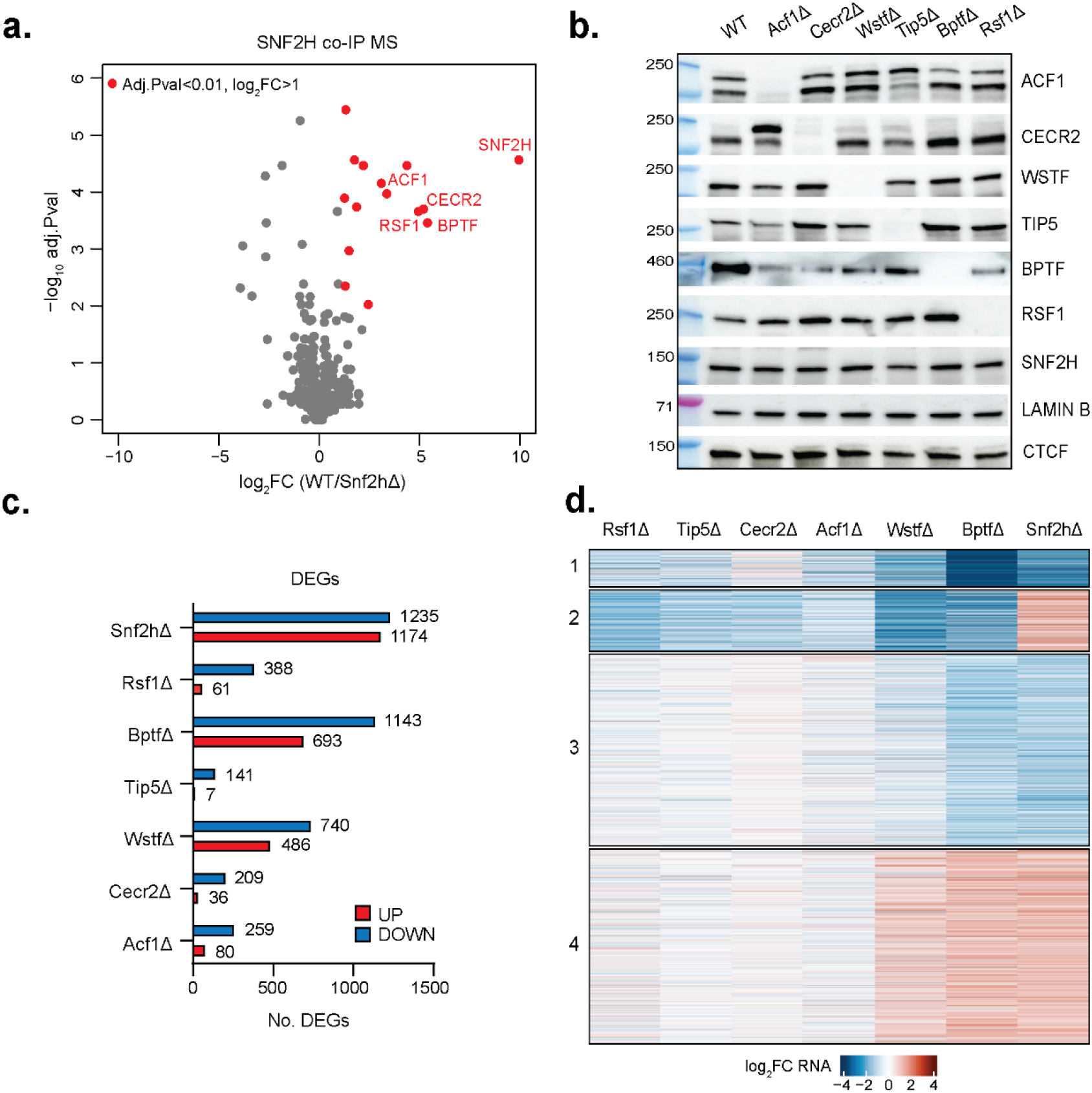
Comprehensive deletion and transcriptome analysis of ISWI subunits in mouse embryonic stem cells. **a.** Mass Spectrometry quantification of proteins co-immunoprecipitated using an anti-SNF2H antibody in wild-type (WT) and Snf2hΔ cells. Highlighted in red are proteins with Benjamini–Hochberg-adjusted two-sided eBayes p-value <0.01 and log_2_ fold-change (WT/ Snf2hΔ) >1. Only names of ISWI subunits are shown. **b.** Western blot for ISWI subunits and controls (LAMIN B, CTCF) in all deletion cell lines and WT control. **c.** Number of differentially expressed genes (DEGs, see Methods) upon deletion of ISWI subunits, upregulated in red and downregulated in blue. **d.** Heatmap of DEGs shown as log_2_ fold-change in respect to parental cell line control. DEGs are clustered based on expression changes, cluster numbers indicated on the left.

Having established the set of SNF2H containing complexes present in mESCs, we generated isogenic loss-of-function mutants using CRISPR/Cas9. More specifically we targeted the main six non-catalytic subunits which distinguish mammalian ISWI subcomplexes, namely *Wstf/Baz1b* for WICH (WstfΔ), *Baz2a/Tip5* for NoRC (Tip5Δ), *Acf1/Baz1a* for the ACF and CHRAC complexes (Acf1Δ), *Rsf1* for RSF (Rsf1Δ), *Bptf* for NURF (BptfΔ), and *Cecr2* for CERF (Cecr2Δ). Genetic deletion was confirmed at the level of DNA sequence and protein expression. All deletion cell lines show absence of detectable protein on western blot, display self-renewal, and express pluripotency markers at levels comparable to wild-type cells (Fig. 1b and Extended data Fig. 1b). First, we asked whether deletion of one subunit would affect the formation of other subcomplexes. More specifically we performed SNF2H co-IP followed by western blot detection of all tested ISWI subunits in each deletion line. As expected, and confirming the genetic deletion, SNF2H co-IP fails to co-immunoprecipitate the deleted subunit in each respective KO cell line (Extended Data Fig. 1a). Importantly, with the above-mentioned exception of BPTF, all other complexes can be detected by western blotting after immunoprecipitation. This suggests that the deletion lines are subcomplex-specific (Extended Data Fig. 1a), and in turn that any molecular phenotype should be a consequence of the loss of individual ISWI subcomplexes.

Next, we investigated subunit-specific transcriptional changes by performing RNA-seq in each of the generated mutant cell lines and parental line as control. Here Tip5Δ, Cecr2Δ, Acf1Δ and Rsf1Δ show moderate transcriptional phenotypes with 148, 245, 339, 449 genes misregulated, respectively (Fig. 1c, Extended Data Fig. 1c). This contrasts with the phenotype of the WstfΔ and BptfΔ lines, which display a much higher number of misregulated genes (1226 and 1836, respectively) (Fig. 1c, Extended Data Fig. 1c). Notably, none of the subcomplex deletions fully recapitulates the transcriptional profile resulting from SNF2H deletion (Extended Data Fig. 2a). However, deletion of the NURF component BPTF causes transcriptional changes that substantially overlap with those upon loss of SNF2H (Extended Data Fig. 2a-b). In order to investigate these further, we performed k-means clustering of all genes differentially expressed across all cell lines (Fig 1d). This identifies two large clusters of genes whose expression changes in a similar fashion in both BPTF and SNF2H depleted cells (Fig. 1d, cluster 3 and 4). However, gene ontology analysis on the respective differentially expressed genes shows no clear enriched terms (Extended Data Fig. 3a-b). Transcriptional changes driven by loss of RSF1, CECR2 and TIP5 were highly correlated albeit modest, affecting genes involved in similar biological processes (Fig. 1d, Extended Data Fig. 3b). Taken together, the transcriptional response to subcomplex specific deletions suggests redundancy among several ISWI subcomplexes and points to a larger and distinct role for the NURF component BPTF.

### Individual ISWI subunits do not account for the nucleosomal phenotype caused by loss of catalytic activity

Next, we asked whether individual deletions of ISWI subunits affect chromatin architecture. Absence of SNF2H causes a global reduction in nucleosome phasing, and coinciding increase in nucleosome repeat length (NRL) by ∼9-10bp^14, 15^. To ask if this can be assigned to an individual subcomplex, we performed Micrococcal-Nuclease sequencing (MNase-seq) which revealed that average NRL is largely unaffected in all of our deletion lines (Fig. 2a). Next, we used the same dataset to ask if nucleosomal phasing at regulatory regions such as transcription start sites (TSSs) and distal DNAseI hypersensitive sites (distal DHSs) is impaired by individual subunit deletions. Similarly to loss of catalytic ISWI activity previously shown in Snf2hΔ cells^14^, phasing at TSSs is unaffected by deletion of individual ISWI subcomplexes (Fig. 2b), suggesting either a reduced role for ISWI in the formation and maintenance of nucleosome structure at TSSs in mouse cells, or potentially a high level of redundancy with other chromatin remodeling complexes such as SWI/SNF. At distal DHSs, deletion of ISWI subunits had similarly little to no effect, with the exception of BPTF (Fig. 2c). Loss of this NURF specific subunit causes increased nucleosomal signal over distal DHSs and coinciding reduction in phasing in the flanking regions (Fig. 2c, Extended data Fig. 4a).

**Figure 2.**
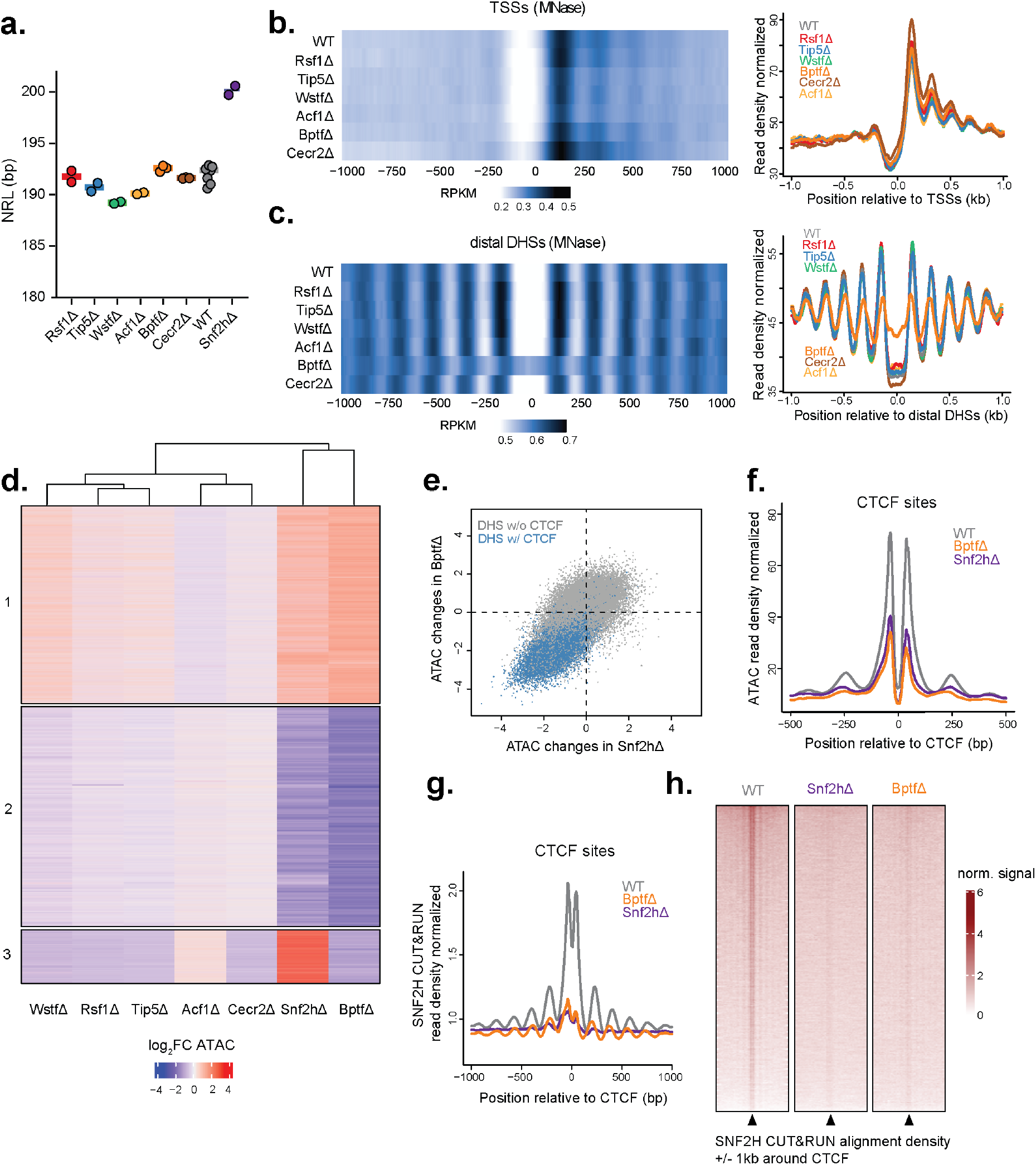
Genome-wide nucleosome position and accessibility profiling reveals sub-complex specific chromatin functions. **a.** Average nucleosome repeat length (NRL) for each deletion cell line and parental line control, as measured by MNase-seq. Shaded bar represents the median. **b-c.** Average nucleosomal profile at transcription start sites (TSSs, b.) or distal DNAseI hypersensitive sites (distal DHSs, c.) shown as heatmap (left) and metaplot (right). **d.** Heatmap displaying log_2_ fold-change of ATAC-seq signal at differentially accessible regions for each deletion cell line in respect to parental line control. Regions are clustered based on accessibility changes and cluster numbers are reported on the left. **e.** Quantitative comparison of chromatin accessibility changes (log_2_ fold-change) driven by genetic deletion of SNF2H (x-axis) and BPTF (y-axis) at DNAseI hypersensitive regions (DHSs). DHSs containing a CTCF motif are highlighted in blue. **f.** Average ATAC-seq signal at bound CTCF motifs in wild-type control (gray), BptfΔ (orange) and Snf2hΔ (purple) cells. **g.** Average SNF2H CUT&RUN signal at bound CTCF motifs (as in **f**) in wild-type control (gray), BptfΔ (orange) and Snf2hΔ (purple) cells. **h.** SNF2H CUT&RUN alignment densities in WT, Snf2hΔ and BptfΔ cells, centered on CTCF bound motifs (black arrow).

Since a large fraction of distal DHSs in mammalian genomes are sites of CTCF binding, we repeated this analysis with a focus on CTCF bound sites, as determined by ChIP-seq. This revealed that the reduction of nucleosomal phasing observed only in BptfΔ cells is indeed concentrated at bound CTCF sites (Extended Data Fig. 4b-c), mirroring our previous observation upon loss of SNF2H^14^.

Next, we asked how the regulatory landscape (as defined by open chromatin) is affected by deletion of ISWI subunits and how close the resulting accessibility profiles are to those occurring upon loss of SNF2H. To this end, we performed ATAC-seq (assay for transposase-accessible chromatin followed by sequencing) in our set of deletion lines and respective parental wild-type controls. We used k-means clustering to categorize open chromatin regions according to their response upon deletion of ISWI subunits. This revealed that the chromatin accessibility landscape is largely unaltered by deletion of CECR2, ACF1, RSF1 and TIP5, in line with the limited transcriptional response in these cells (Fig. 2d). Again, the BPTF mutant line stands out as it displays major changes in genome-wide chromatin accessibility, which extensively overlap with the changes driven by deletion of the catalytic subunit SNF2H (Fig. 2d, Extended Data Fig. 4d).

Further analysis of the distribution of chromatin marks and chromatin binding factors revealed that regions characterized by loss of accessibility in both BptfΔ and Snf2hΔ (cluster 2 in Fig. 2d) are highly enriched for CTCF binding, compared to any other clusters or any other functional annotation (Extended Data Fig. 4e). In fact, motif enrichment analysis revealed very high enrichment for CTCF motifs in regions allocated to cluster 2, suggesting that loss of accessibility driven by deletion of either BPTF or SNF2H is mostly localized at CTCF motifs (Fig. 2e-f, Extended Data Fig. 4f). This characteristic, which mirrors the phenotype of Snf2hΔ cells, is limited to the BPTF deletion and it is not observed in any other deletion line (Extended Data Fig. 4g). Taken together, genome-wide accessibility maps reveal only modest effects upon deletion of NoRC, RSF, CERF and ACF complexes in mESCs while deletion of BPTF causes large changes at CTCF sites that resemble the phenotype observed in Snf2hΔ cells.

To test if the specific impact on CTCF sites reflects local recruitment of remodeler activity via BPTF, we investigated whether ISWI localizes to CTCF sites in a BPTF-dependent manner. More specifically, we profiled SNF2H genomic binding in the presence and absence of BPTF using CUT&RUN^17^ including SNF2H depleted cells as negative control. This revealed that in both wild-type and BptfΔ cells, SNF2H localizes at distal accessible regions represented by distal DNAseI hypersensitive sites (Extended Data Fig. 5a), highlighting that BPTF depletion does not globally alter the ATPase localization at distal regulatory sites. However, in the absence of BPTF, SNF2H signal is lost specifically at sites occupied by CTCF in wild-type cells (Fig. 2g-h, Extended Data Fig. 5b-d). Interestingly, SNF2H localization decreases, although to a lower extent, also at CTCF sites which retain accessibility in absence of BPTF (Extended Data Fig. 5e). Altogether, the reduction in SNF2H localization upon BPTF depletion is in line with the observed accessibility changes and suggests that the NURF remodeling complex mediates SNF2H localization and function at CTCF sites.

### NURF-component BPTF mediates accessibility upon CTCF binding

Loss of accessibility upon remodeler deletions has thus far coincided with loss of TF binding^14, 18, 19^. This has also been observed for CTCF upon deletion of SNF2H and can thus be expected to be the case upon deletion of BPTF.

To test this, we performed chromatin-immunoprecipitation followed by sequencing (ChIP-seq) for CTCF in wild-type and BptfΔ cells. In contrast to expectations, most sites that show drastic reduction in accessibility remain either bound by CTCF or display a minor reduction in binding, which contrasts with strong reduction in binding in the Snf2hΔ (Extended Data Fig. 6a). This is evident when looking at the average CTCF ChIP-seq signal through all bound sites which shows stronger loss in SNF2H versus BPTF depleted cells (Fig. 3a); but also, at individual sites where CTCF binding is consistently lost upon SNF2H deletion while the response to BPTF deletion is weaker or absent (Fig. 3b). This indicates that genetic deletion of the NURF component BPTF results in a state distinct from complete loss of ISWI function, where CTCF binding can persist, while chromatin opening at the same sites is strongly impaired.

**Figure 3.**
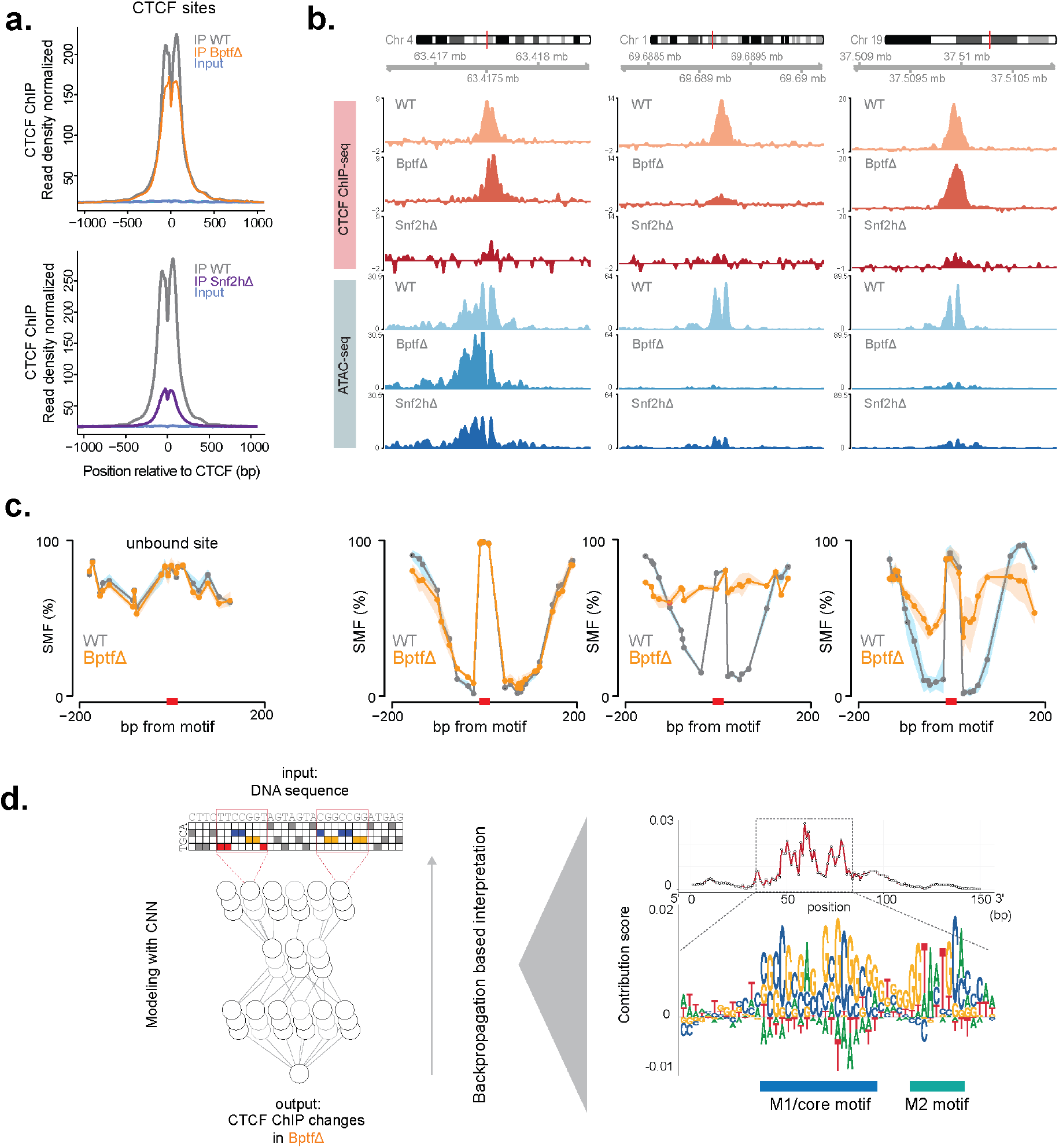
CTCF binding at strong motifs largely persists in the absence of BPTF despite loss of accessibility. **a.** Average CTCF ChIP-seq signal at bound CTCF sites in BptfΔ (orange) and parental ES cells (gray) (top panel). Same analysis in Snf2hΔ (purple) and parental ES cells (gray) (data from ref.^14^) (bottom panel). Inputs are shown as control (blue). **b.** Representative genomic loci illustrating changes in CTCF binding (ChIP-seq - shades of red) and chromatin accessibility (ATAC-seq - shades of blue) in BptfΔ, Snf2hΔ and parental ES cells. **c.** Average single-molecule footprinting (SMF) signal at an unbound site (left) and at the same sites (as in **b,** right), in BPTF deleted cells (orange) and wild-type control (gray). Shaded line represents standard deviation. **d.** A convolutional neural network-based model used to predict changes in CTCF binding in BptfΔ cells. Influence of particular nucleotides is shown as average contribution scores highlighting the role of an extended CTCF motif (M1 and M2) in retaining binding in absence of BPTF.

### Single molecule footprinting confirms persistent CTCF binding upon BPTF deletion

Thus far we relied on enrichment based-methods (ChIP-seq and ATAC-seq) to identify the role of BPTF at sites of CTCF binding. This leaves the possibility that the observed disconnect between binding and accessibility could be a function of differential sensitivities of each assay to local changes. Experimentally testing of this possibility requires an approach that does not rely on enrichment and that, ideally, detects both TF binding and nucleosomal organization simultaneously. To this end we performed Single-Molecule Footprinting (SMF or NOMe-seq), a technique that utilizes exogenous enzymatic methylation of cytosines in the GpC context (which are normally endogenously unmethylated), which is blocked by the presence of nucleosomes or DNA-bound factors, allowing single-molecule and base-resolution footprinting on chromatin^20^. More specifically we performed amplicon-based SMF in wild-type and BptfΔ mESCs. The regions tested are representative of sites that upon BPTF deletion either 1) maintain both CTCF binding and chromatin accessibility, 2) show loss of both features, or 3) maintain CTCF binding despite loss of accessibility (as profiled by ChIP-seq and ATAC seq, Fig. 3b from left to right). In addition, we included an unbound CTCF site as baseline reference. The resulting footprinting patterns at the tested locations reveal that sites with unchanged CTCF ChIP-seq and ATAC-seq signal upon BPTF deletion also show identical footprinting as the wild-type cells (Fig. 3c second panel from the left). Loss of CTCF binding in BptfΔ is paired with flattening of average footprinting signal, concordant with signal coming from unbound sites (Fig. 3c third and first panel from the left respectively). Crucially, sites that maintain CTCF binding but loose accessibility, maintain a clear footprint over the CTCF motif upon BPTF deletion, with same occupancy as in wild-type cells. However flanking regions show reduced methylation signal and thus impaired accessibility for the footprinting enzyme. This mirrors the accessibility reduction observed in ATAC seq (Fig. 3c fourth panel from the left).

Finally, we reasoned that reduced accessibility proximal to bound CTCF sites could reflect increased nucleosome occupancy due to lack of NURF remodeling. If indeed the case, this should result in higher abundance of longer MNase fragments over bound CTCF sites in BPTF depleted cells compared to wild-type control. Indeed, analysis of the relative enrichment of MNase-seq reads as a function of fragment size shows a clear accumulation of longer (> 200bp) fragments that span the bound motifs but only in the absence of BPTF (Extended Data Fig. 6b). Taken together, these experiments indicate that without the NURF component BPTF, binding of CTCF largely persists, but it is not accompanied by canonical opening and creation of hyper-accessible chromatin.

### An extended and conserved CTCF motif is a feature of sites with persistent binding in absence of NURF

Next, we wanted to investigate why different sites show different CTCF binding response in the absence of NURF remodeling. With this aim, we asked how persistent binding of CTCF upon NURF depletion relates to initial binding strength or DNA sequence. Interestingly, CTCF ChIP-seq signal in wild-type cells did not correlate with loss of CTCF binding upon BPTF depletion (Extended Data Fig. 7a). Motif strength, however, showed a weak yet noticeable trend, in that CTCF sites with higher motif scores tend to retain more CTCF binding when BPTF is absent (Extended data Fig. 7b). To go beyond prior motif definitions, we explored how persistent CTCF binding relates to sequence features using a convolutional neural network approach (Fig. 3d). We took 150bp of DNA sequence around CTCF motifs as independent variable and the response in CTCF ChIP-seq signal in BptfΔ cells as the dependent variable. The employed network architecture was similar to the recently published DeepSTARR network, which has been used to map DNA sequence to tissue-specific enhancer activity (Methods and ref.^21^). This model shows a relatively high level of correlation when used to predict CTCF ChIP-response upon BPTF depletion in held-out test chromosomes (Extended Data Fig. 7c-d). In line with the previous observation that sites with higher motif score are more likely to preserve CTCF binding, the deep learning approach found that the canonical CTCF motif has predictive value explaining the persistent CTCF binding upon BPTF depletion (Fig. 3d). In addition, this approach also revealed the contribution of an additional 9-nt stretch located ∼21bp from the center of the canonical CTCF motif (Fig. 3d, Extended Data Fig. 7e). This 9-bp motif corresponds to an additional CTCF motif component (“M2 motif”), reported in a minority of CTCF binding events and originally discovered in DNAseI datasets^22, 23^. M2-containing CTCF sites have been previously associated with highly evolutionary conserved binding events which also tend to be less sensitive to CTCF protein knockdown^23^.

Here, the deep learning approach revealed the contribution of both M1 and M2 motifs to the persistent CTCF binding in BPTF depleted cells. Of note, most CTCF sites do not harbor an M2 motif. While this is the case for the entirety of all CTCF sites, we observe that the fraction of sites containing an M2 sequence increases for sites characterized by persistent binding in BptfΔ cells (Extended Data Fig. 7f). Combined, these findings suggest that not only strong canonical motifs but also the presence of additional motif components, likely contributing to protein-DNA affinity, can reduce dependence on remodeling activity for binding.

### Absence of BPTF reduces long-range chromatin interactions

To further explore the relationship between chromatin accessibility and CTCF binding we clustered CTCF sites based on changes in accessibility and binding in BptfΔ cells.

Within each cluster, we then compared the average ATAC-seq and CTCF ChIP-seq responses to the deletion of BPTF and SNF2H (Fig. 4a, Extended Data Fig. 8a-b). In line with the above observations, this analysis illustrates a much stronger reduction of CTCF binding in Snf2hΔ versus BptfΔ cells within all clusters, while the reduction in accessibility is similar in both mutants (Fig. 4a). Notably, the resulting clusters showed distinct patterns of chromatin states as previously defined by the combinations of histone marks and TF binding in mESCs ^24, 25^ (Fig. 4b). This analysis showed that the cluster of sites with limited changes in accessibility in both mutants is highly enriched for promoter and enhancer states (cluster 1 Fig. 4a-b, Fisher’s exact p-value < 2.2e-16, odds ratio=6.1). These CTCF sites are likely embedded within regulatory regions, suggesting that accessibility is maintained by the activity of other TFs.

**Figure 4.**
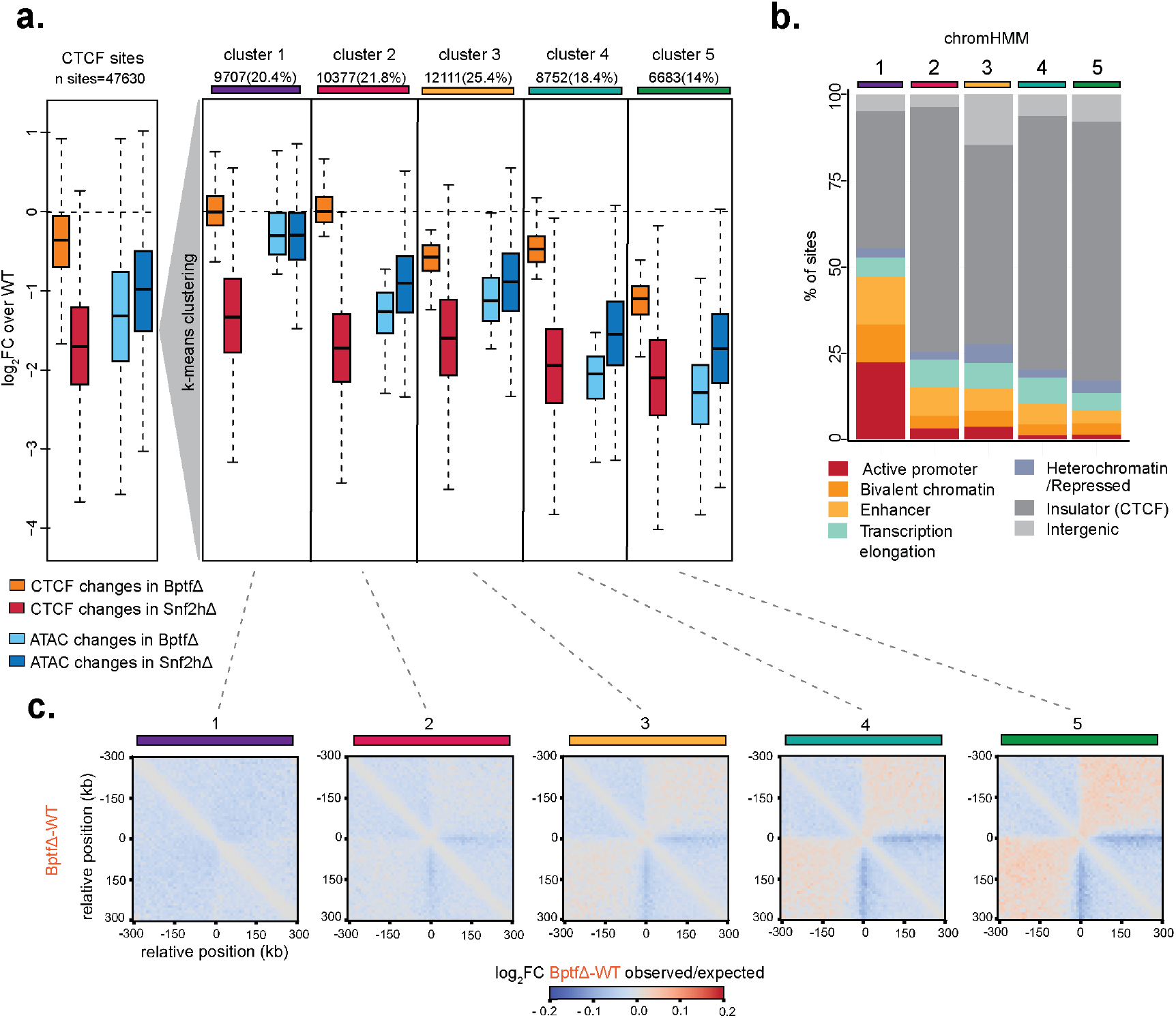
Unsupervised clustering highlights differential responses to absence of BPTF at the level of CTCF binding, chromatin opening and nuclear organization. **a.** Boxplot displaying changes upon BPTF or SNF2H deletion in ChIP-seq (shades of red) and ATAC-seq (shades of blue) signal at all bound CTCF sites, expressed as log_2_ fold-change in respect to wild-type control cells. Measurements are shown for all sites (left), and then separately for clusters 1-5. Central line represents the median to first/third quartile, whiskers to 1.5 multiplied by interquartile range. **b.** Distribution of chromatin states at sites surrounding bound CTCF motifs, as labeled by chromHMM and split by clusters (as in **a**). Cluster numbers reported on top. **c.** Cluster-specific changes in observed/expected interactions at CTCF sites following BPTF deletion, measured using Hi-C. Cluster numbers reported on top.

Next, we asked if the insulator function at CTCF sites is affected upon BPTF deletion, as we have previously observed for the SNF2H mutant^14^. Hi-C experiments indeed reveal a global reduction in long range interactions in BptfΔ (Extended Data Fig. 8c). Dividing the changes in contact frequency by the five clusters as defined (Fig. 4c, Extended Data Fig. 8d) suggests that CTCF sites with increased loss of accessibility show more significant reduction in long-range interactions. This opens the possibility of changes in the 3D nuclear organization as a function of reduced accessibility.

### BPTF-mediated accessibility around CTCF sites is required for efficient insulation

Given the limited reduction in CTCF binding in the BptfΔ cells, we hypothesized that reduced accessibility around bound CTCF sites could account for the impairment of insulator function. If true, upon deletion of BPTF, there should be many sites with persistent binding but reduced insulation.

To challenge this model, we focused on those 15865 CTCF sites with only very limited variation in binding (+/- 20% variation in ChIP-seq signal in BptfΔ), grouped these according to their changes in accessibility (Fig. 5a, Extended Data Fig. 9a) and calculated 3D contact frequencies within these groups (Fig. 5b, Extended Data Fig. 9b). This reveals that stronger loss of accessibility is paired with increased loss of insulator function in line with the original hypothesis. Next, we tested if this impairment in insulator function coincides with reduced interaction with cohesin, measured by ChIP-seq for its component RAD21. Indeed, RAD21 shows reduced binding in the BPTF mutant, scaling with the reduced accessibility, even at sites with little to no change in CTCF binding (Fig. 5c top panels). This behavior is not restricted to cohesin but can be similarly observed for its release factor WAPL, which shows a comparable decrease (Fig 5c bottom panels).

**Figure 5.**
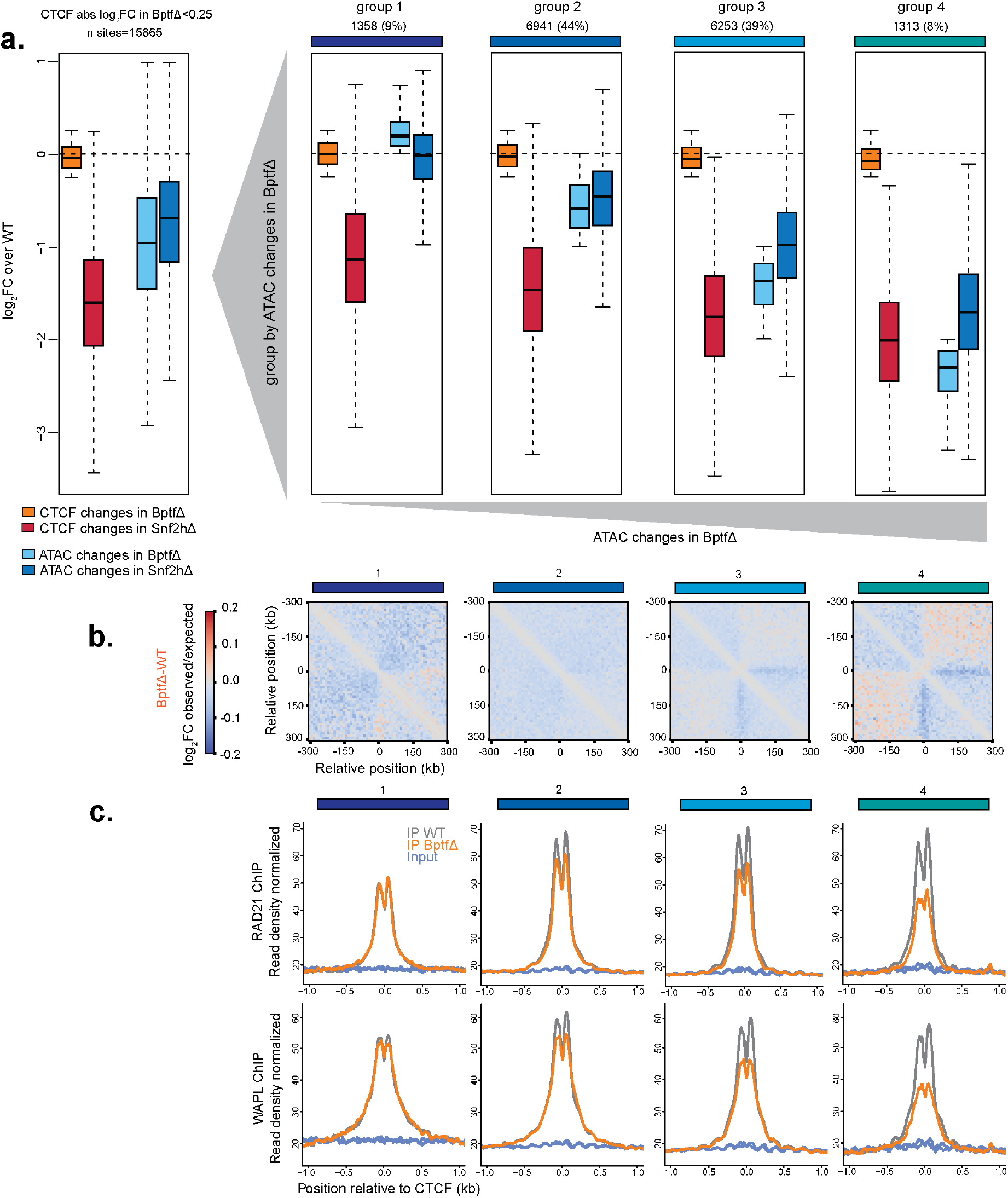
BPTF-dependent accessibility impacts nuclear organization, localization of cohesin and cohesin-release factors despite persistent CTCF binding. **a.** Boxplot summarizing changes in ChIP-seq (shades of red) and ATAC-seq (shades of blue) signal for CTCF sites that retain binding upon BPTF deletion, expressed as log_2_ fold-change in respect to wild-type control cells. Measurements are shown for all sites that retain binding (left), and divided into groups 1-4 based on their accessibility changes upon loss of BPTF. **b-c.** Changes in observed/expected interactions upon BPTF deletion (b) and average ChIP-seq signal for RAD21 and WAPL (c), at CTCF sites that retain binding, grouped by changes in chromatin accessibility (as in **a**).

We conclude that proper insulator function at CTCF sites requires not only the presence of bound CTCF but also chromatin opening mediated by NURF. The BPTF deletion phenotype further reveals that CTCF binding does not necessarily require chromatin opening and that CTCF binding is not sufficient for chromatin opening. These results separate local CTCF binding from insulator function in nuclear organization.

## Discussion

Our systematic analysis reveals a specific function for the NURF component BPTF in creating accessibility at CTCF sites, which we show to be critical for insulator function but not CTCF binding. Furthermore, we show that no single ISWI subcomplex accounts for the changes in nucleosome repeat length observed when depleting the catalytic subunit SNF2H^14, 15^. This argues for a redundancy by at least two subcomplexes in regulating nucleosome distance and thus in providing the general nucleosome mobility catalyzed by ISWI. This finding is compatible with the fact that ACF, RSF and WICH mammalian complexes are all able to space nucleosomes in vitro^6–8^.

The observed phenotype upon deleting the NURF subunit BPTF is unexpected as it recapitulates only a specific part of the SNF2H loss-of-function phenotype previously observed at CTCF bound sites^14, 15, 26^, in that the accessory subunit BPTF is required for maintaining these sites in an open chromatin state. This argues for a specific and local function for NURF catalyzed by SNF2H, which we show localizes to CTCF sites in a BPTF-dependent manner in line with their biochemical interaction. In contrast to the phenotype observed upon SNF2H deletion, CTCF binding unexpectedly remains largely retained in the absence of BPTF. This genome-wide observation upon a stable genetic deletion is compatible with previous findings at a subset of individual CTCF sites^27^ or reported nucleosomal profiles upon siRNA depletion of different ISWI accessory subunits^26^, respectively in mouse and human cellular models.

CTCF affinity for its motif seems to be crucial in retaining binding in absence of NURF, as shown by a dependency of residual binding on motif score, including an additional motif component previously described as the “M2 motif” that extends the canonical M1 motif of CTCF^23^. This additional sequence component has been proposed to interact with the CTCF zinc fingers 9 to 11^28, 29^. Moreover, CTCF sites containing this motif have been reported to be more resistant to CTCF knock-down in human cells, suggesting higher affinity at these regions^23^. This supports a model where higher motif affinity leads to more stable CTCF binding and a reduced requirement for general ISWI activity, which remains to be tested. The difference in phenotype between SNF2H and BPTF deletions argues that ISWI activity other than NURF enables CTCF binding. Since preferential SNF2H occupancy at CTCF sites requires NURF we speculate that unspecific nucleosome mobilization by other ISWI subcomplexes can create binding opportunities.

Several scenarios have been proposed about how TF binding and chromatin opening are linked^30–32^. These involve unspecific remodeler activity that is stabilized by TF binding or TF-dependent recruitment of remodeler activity. In the case of CTCF, our findings let us propose a multistep model where unspecific nucleosomal mobility mediated by ISWI enables CTCF binding followed by specific recruitment of NURF, which leads to chromatin opening.

The separation of binding from chromatin opening by deleting BPTF enabled us to test potential functional consequences and revealed that CTCF binding alone is insufficient for proper insulation which appears to require chromatin opening. Since this coincides with reduced presence of cohesin, it is tempting to speculate that chromatin opening enables stable recruitment of the cohesin factors and consequent long range contact formation. This model might also apply to *Drosophila* where NURF-301 (BPTF) contributes to insulation^33, 34^.

Combined, this suggests an evolutionary conserved function of NURF at insulators and argues that, at least in the studied system, NURF-mediated chromatin remodeling is necessary for CTCF function as insulator post chromatin binding.

Since its first description, open chromatin has become an established hallmark of active regulatory regions^35^. It is so canonical that it is successfully used for their comprehensive genome-wide detection in any given cell type^30, 32^. Chromatin opening itself depends on binding of TFs that are chromatin insensitive, also referred to as pioneer factors, and further thought to enable binding for chromatin sensitive factors^31^. Several studies have shown that chromatin opening relies on the activity of very different remodeler families including ISWI and SWI/SNF and occurs in a TF specific fashion^14, 18, 19, 36^. In the reported cases removal or inhibition of remodeler activity caused reduced binding and coinciding reduced accessibility around binding sites of selected TFs. However, we are not aware of an example where absence of a cofactor causes loss of chromatin opening yet, largely persistent TF binding as we show here for CTCF in absence of BPTF.

The fact that chromatin opening is critical for insulator function independently of CTCF DNA binding adds a previously underappreciated variable to the set of requirements for proper nuclear organization. More generally, it suggests a model where local chromatin opening enables or assists cofactor interaction for TFs, potentially creating the necessary space and flexibility for proper regulatory function. If true, similar phenotypes, might be identified with other TFs either by mutating cofactors as presented here or by mutating cofactor interacting domains of TFs. Ultimately, this should shed further light on how chromatin structure and nucleosome mobility influence TF function beyond DNA binding.

## Methods

### Cell culture

Wild-type (WT) mESCs and derived genetically deleted lines of 129S6/SvEvTac background were maintained as previously described^19^. Briefly, cells were maintained in Dulbecco’s modified Eagle’s medium (Invitrogen), supplemented with 15% fetal calf serum (Invitrogen), GlutaMAX Supplement (Gibco) and nonessential amino acids (Gibco), β-mercaptoethanol (Sigma) and leukemia inhibitory factor (produced in house). Cells were grown on plates coated with 0.2% gelatin (Sigma).

### Cell line generation and maintenance

All deleted cell lines were generated starting from mESC lines of 129S6/SvEvTac background using the CRISPR/Cas9 protocols previously described, with modifications^37^. Briefly, mouse ES cells were co-transfected (Lipofectamine 3000, Thermo Fisher Scientific) with vectors expressing Cas9 and sgRNAs targeting the gene of interest. Puromycin selection (2 μg/ml) was carried out one day after transfection for 24h. Following a 2-day recovery, individual colonies were expanded and genotyped by western blot and DNA sequencing. For each gene, multiple deleted clones were isolated from separate transfections with separate gRNAs to assess off-target and clonal effects. Sequence of gRNAs used for this study are reported in the table below:

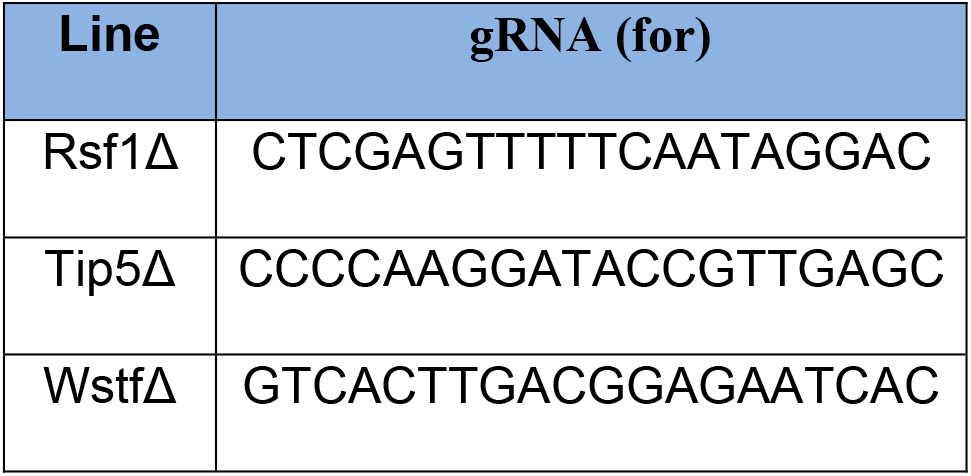
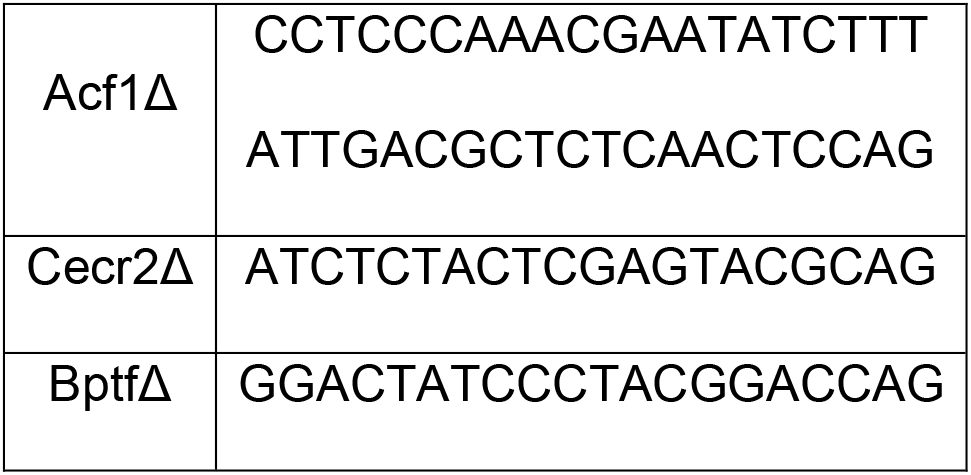

### Nuclear protein extraction for western blotting and co-immunoprecipitation

For nuclear cell lysis 5×10^6 cells per condition were resuspended in 1mL of Lysis Buffer + protease inhibitor (cOmplete, Roche) (10mM Tris-HCl pH 7.4, 10mM NaCl, 3mM MgCl2, 0.1mM EDTA, 0.5% NP40) and incubated on ice for 10 min. Samples were then centrifuged for 5 min at 3000rpm at 4°C and resuspended in 250µl of Wash Buffer + protease inhibitor (10mM Tris-HCl pH 7.4, 10mM NaCl, 3mM MgCl2, 0.1mM EDTA). Samples were then directly centrifuged for 5min at 3000rpm 4°C. Resulting nuclei were resuspended in 250 µl RIPA-buffer + protease inhibitor (50mM Tris-HCl pH 8.0, 150mM NaCl, 1% NP40, 1% Na-deoxycholate, 0.1%SDS, 1mM DTT), mixed briefly through vortexing and incubated on ice for 30 min. After this time, samples were sonicated twice (7 cycles 30s on, 30s off) with a Bioruptor Plus sonicator (Diagenode). In between the two sonication cycles samples were incubated at 12°C with 0.8ul of Benzonase (Sigma). Finally, lysates were centrifuged at 4°C for 15 min at 13.000rpm and resulting supernatant was used for SDS-PAGE and western blotting directly or for co-immunoprecipitation followed by SDS-PAGE and western blotting.

### Co-immunoprecipitation followed by western blotting

For co-immunoprecipitation, protein G Dynabeads magnetic beads (Thermo Fisher Scientific) were washed twice and resuspended in their original volume with RIPA buffer diluted 1:1 in dilution buffer (10mM Tris-HCl pH 7.5, 150mM NaCl, 0.5mM EDTA, 1mM DTT, protease inhibitor). Nuclear extracts for the IP were obtained as described in the previous section. For preclearing, 15ul of beads were added to nuclear extracts and incubated for 30 min. at 4°C with overhead rotation. 5% of the precleared lysate was used as input control. To the rest of the precleared lysates 5ul of anti-SNF2H-antibody were added and samples were let overnight at 4°C with rotation. The following day, 25ul of prewashed magnetic beads were added and samples were let 1h at 4°C with rotation. After this time, beads were washed 3 times with 500ul IP-wash buffer (20mM TRIS-HCl pH 8.0, 150mM NaCl, 1mM EDTA, 1.5mM MgCl2, 0.5%NP40, 1mM DTT, protease inhibitor). Beads were then transferred to a new tube and washed again twice before being resuspended in appropriate volume of Gel-loading buffer (1:1 mix of RIPA buffer and 5x Laemmli-buffer/5% β-mercaptoethanol) for SDS-PAGE and western blotting.

### Co-immunoprecipitation followed by mass spectrometry

In brief, the day before the experiment WT and Snf2hΔ mES cells were seeded into 15 cm plate (1×10^7 cells/plate). For each condition three independent biological replicates were prepared. For the lysis, 1.5×10^7 cells per sample were washed in PBS and resuspended in 1mL of hypotonic solution (20 mM Tris-HCl pH 7.4, 10mM NaCl, 3mM MgCl2) and incubated on ice for 5 min. After incubation, NP40 was added to a final concentration of 0.1% ; samples were mixed gently, left on ice for 5 min and centrifuged for 5 min at 500g. Resulting pellets were resuspended in 900ul of B150AG buffer (10mM Tris-HCl pH 7.5, 2mM MgCl2, 150mM NaCl, 0.5% Triton X-100, 50mM L-Arg [Sigma A5006], 50mM L-Glu [SigmaG1251] + protease inhibitor) and mixed by vortexing. After that samples were centrifuged for 5 min at 4°C at maximum speed. Supernatants were then transferred to new tubes for immunoprecipitation.

In brief, for each IP 25µL of protein G Dynabeads magnetic beads (Thermo Fisher Scientific) were washed twice with 1mL of B150 buffer (10mM TrisHCl pH 7.5, 2mM MgCl2, 150mM NaCl, 0.5% Triton X-100). After washing, beads were resuspended in 100µL of B150 buffer + protease inhibitor and incubated with lysates for 1h at 4°C. Precleared lysates were moved to fresh tubes and incubated at 4°C overnight with 5ug of a-Snf2h antibody. The day after, 25µL of beads per IP were washed as described above and incubated with samples at 4°C for 4h. After incubation, beads were washed 3 times with B150 buffer, resuspended in 250µL of B150nd buffer (10mM TrisHCl pH 7.5, 2mM MgCl2, 150mM NaCl) and moved to clean tubes. Then beads were again washed on magnet with 1mL of B150nd buffer, supernatant was removed, and beads were centrifuged for 30s at maximum speed.

Protein digestion was carried out as previously described^38^. In brief, beads were resuspended in 5µl Digestion buffer (3M guanidinium hydrochloride, 20mM EPPS pH 8.5, 10mM chloroacetamide, 5mM TCEP) and digested with 1µl of 0.2µg/µl Lys-C at room temperature for 4h. 17uL 50mM HEPES pH 8.5 were added to the beads, followed by the addition of 1µl 0.2µg/µl trypsin. Beads were then incubated at 37°C overnight. The day after, another 1µl of 0.2µg/µl trypsin was added and samples were digested for an additional 5h. Samples were acidified by adding 1uL of 20% TFA and sonicated in an ultrasound bath. Peptides were analyzed by LC–MS/MS on an EASY-nLC 1000 (Thermo Scientific) using a two-column set-up. The peptides were applied onto a peptide μPAC™ trapping column in 0.1% formic acid, 2% acetonitrile in H2O at a constant flow rate of 5μl/min. Using a flow rate of 500 nl/min, peptides were separated at room temperature with a linear gradient of 3%–6% buffer B in buffer A in min followed by a linear increase from 6 to 22% in 55 min, 22%–40% in 4 min, 40%– 80% in 1 min, and the column was finally washed for 13 min at 80% buffer B in buffer A (buffer A: 0.1% formic acid; buffer B: 0.1% formic acid in acetonitrile) on a 50cm μPAC™ column (PharmaFluidics) mounted on an EASY-Spray™ source (Thermo Scientific) connected to an Orbitrap Fusion LUMOS (Thermo Scientific). The data were acquired using 120,000 resolution for the peptide measurements in the Orbitrap and a top T (3 s) method with HCD fragmentation for each precursor and fragment measurement in the ion trap according to the recommendation of the manufacturer (Thermo Scientific).

### Antibodies

| Antibody | Application | Vendor | Cat. Number |
| --- | --- | --- | --- |
| Snf2h | WB, CUT&RUN | Abcam | ab72499 |
| Snf2h | IP | Abcam | ab3749 |
| CTCF | ChIP-seq | Santa Cruz | C-20X |
| CTCF | WB | Abcam | ab128873 |
| Rad21 | ChIP-seq | Abcam | ab992 |
| Wapl | ChIP-seq | Proteintech | 16370-1-AP |
| IgG | CUT&RUN | Sigma | M7023 |
| Acf1 | WB | Bethyl<br>Laboratories | A301-318A |
| Cecr2 | WB | LSBio | LSC-496852 |
| Wstf | WB | Abcam | ab 51256 |
| Tip5 | WB | Diagenode | C15310090 |
| Bptf | WB | Millipore | ABE24 |
| Rsf1 | WB | Abcam | ab 109002 |
| Lamin B | WB | Abcam | ab 16048 |

### RNA-seq

Total RNA for RNA-seq was purified using RNeasy Mini Kit (QIAGEN), and any residual genomic DNA was removed using a DNA-free DNA Removal Kit (Invitrogen). Sequencing libraries were prepared from purified RNA for two or three biological replicates using TruSeq RNA Library Prep kit v2 (Illumina). Libraries were sequenced on the Illumina HiSeq 2500 or Illumina NovaSeq.

### ATAC-seq

ATAC-seq was performed according to previously described protocols^19^. Briefly, 50,000 cells were washed with cold phosphate-buffered saline and resuspended in 50 μl of lysis buffer (10 mM Tris-HCl pH 7.4, 10 mM NaCl and 3 mM MgCl_2_, 0.1% NP40, 0.1% Tween-20 and 0.01% digitonin) and incubated on ice for 3 minutes to extract the nuclei. After lysis, 1 ml of wash buffer (10 mM Tris-HCl pH 7.4, 10 mM NaCl and 3 mM MgCl_2_, 0.1% NP-40) was added, and the tubes were inverted to mix. The nuclei were cold centrifuged at 500*g* for 10 min. The nuclei pellet was incubated in 50 μl of transposition reaction buffer (25 μl 2 × TD buffer, 2.5 μl transposase (100 nM final), 16.5 μl PBS, 0.5 μl 1% digitonin, 0.5 μl 10% Tween-20 and 5 μl water) for 30 min at 37 °C in an orbital shaker at 750 rpm. The DNA was purified using the MinElute PCR Purification Kit (QIAGEN). The eluted transposed DNA was submitted to PCR using Q5 High-Fidelity Polymerase (New England Biolabs). DNA was amplified with 7 cycles of PCR. The libraries were sequenced on the Illumina NextSeq platform at 41 bp paired-end. All ATAC-seq experiments were performed in at least two independent biological replicates per condition.

### MNase-seq

MNase-seq was performed as previously described^19^, in brief: cells were resuspended in 1mL of Buffer 1 (0.3M Sucrose, 15mM Tris pH 7.5, 60mM KCl, 15mM NaCl, 5mM MgCl2, 2mM EDTA, 0.5mM DTT, 1x PIC, 0.2mM spermine, 1mM spermidine) w/ detergent (Buffer 1 + 0.02% NP-40) and incubated on ice for 5min. Nuclei were then pelleted at 300g for 5min at 4°C. Nuclei were gently resuspended in 1mL of Buffer 2 (0.3M Sucrose, 15mM Tris pH 7.5, 60mM KCl, 15mM NaCl, 5mM MgCl2, 0.5mM DTT*, 1 x PIC, 0.2mM spermine, 1mM spermidine). Nuclei were then pelleted for 5 min at 300g at 4°C. Pellets were resuspended in 400 μl MNase buffer (0.3M Sucrose, 50mM Tris pH 7.5, 4mM MgCl2, 1mM CaCl2, 1 x PIC). 5U of MNase S7 micrococcal nuclease (Roche) were added. Nuclei were then incubated for 30min at 37°C. Reaction was stopped by adding EDTA to a final concentration of 5 mM. SDS (to a final concentration 1%) and Proteinase K (200 μg/mL) were then added to the samples, followed by incubation at 55°C for 1h with shaking. MNase-digested DNA was purified using AMPure XP beads. 500 ng of purified DNA was then used for library preparation with NEB Next Ultra Library Preparation Kit (New England Biolabs), using five PCR cycles. Libraries were sequenced on the Illumina NextSeq 500 (41 bp paired-end) or Illumina NovaSeq. All MNase-seq experiments were performed in at least two independent biological replicates per condition.

### Single-molecule Footprinting and amplicon Bisulfite sequencing

Single-molecule footprinting was carried out as previously described^39^ for wild-type and BPTFΔ cells in three independent biological replicates each. For each sample, 2μg of methylated DNA was used for bisulfite conversion using EZ DNA methylation-gold kit (Zymo). Converted DNA was used as a substrate for PCR amplification of the below endogenous CTCF sites using bisulfite-compatible primers and KAPA HiFi Uracil+ (Roche) ([95 °C, 4 min] × 1; [98 °C, 20 s; 60 °C, 15 s; 72 °C, 20 s] × 35, [75 °C, min] × 1; 4 °C, hold):

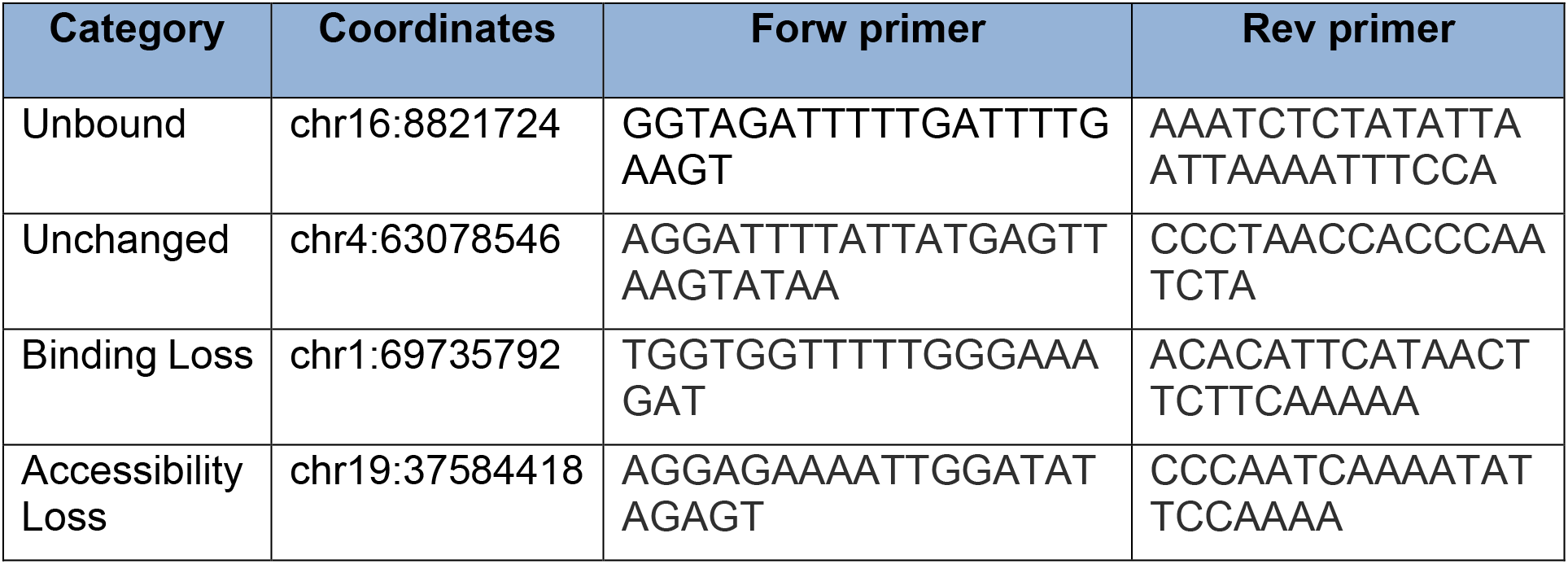

Amplicons were purified using AMPure XP beads (Beckman Coulter), pooled by sample, and used for library preparation using NEBNext Ultra Library Preparation Kit (New England Biolabs) and sequenced on an Illumina MiSeq (250bp paired-end).

### CUT&RUN

CUT&RUN was performed following the EpiCypher manufacturer’s protocol version 1.5.2 (https://www.epicypher.com/technologies/cutana/cut-and-run) with some modifications. For all conditions CUT&RUN was performed with both Snf2h and IgG antibody as control in two independent biological replicates. In brief, the day before the experiment WT, Snf2hΔ, and BptfΔ mES cells were seeded in 6-well plates. The day after 10µL/sample of Concanavalin A beads (Concanavalin A magnetic beads, Bangs Laboratories) were washed twice with Bead Activation Buffer (20mM HEPES pH 7.9, 10mM KCl, 1mM CaCl2, 1mM MnCl2) and resuspended in 10uL of the same buffer. For each sample 0.5 x 10^6 cells were washed in PBS and centrifuged at room temperature at 600g for 3 min. Cells were then washed twice in 100µL of Wash buffer (20mM HEPES pH7.5, 150mM NaCl, 0.5mM Spermidine, protease inhibitor) and resuspended in 100uL of the same buffer. Afterwards, cells were aliquoted in 8-strip tubes containing 10 µL of activated beads, mixed with gentle vortexing and incubated at room temperature for 10 min. After this time, supernatant was removed and beads were gently resuspended in 50 µL of cold Antibody buffer (Wash Buffer + 0.001% Digitonin + 2mM EDTA). 0.5µL of antibody were added to each sample and let overnight at 4°C on a nutator. The following day, beads were washed twice using 250µL of cold Digitonin Buffer (Wash buffer + 0.001% Digitonin) and gently resuspended in 50µL of the same buffer. 2.5µL of CUTANA pAG-MNase (20X pAG-MNase, Epicypher) were added to each of the samples, which were then gently mixed and left 10min at room temperature. After this time, 250µL of cold Digitonin buffer were added directly to samples. The previous step was repeated for two washes then samples were resuspended in 50µL of Digitonin buffer. To start the digestion, 2uL of 50mM CaCl2 were added to the samples which were then gently mixed and let at 4°C for 2h on a nutator. After this time 33µL of Stop Buffer (340 mM NaCl, 20mM EDTA, 4mM EGTA, 50µg/mL RNase A, 50µg/mL Glycogen) were added, samples were then vortexed and incubated at 37°C for 10 min. Samples were moved on magnet and supernatant was transferred to clean 1.5 mL tubes for nucleic acid extraction using the MinElute PCR Purification Kit (Qiagen). Purified DNA was used for library preparation using the NEBNext Ultra Library prep Kit (Illumina) according to manufacturer’s instructions with the following Epicypher manufacturer’s modifications. DNA clean up before PCR amplification was done using 1.1X AMPure XP beads. PCR amplification parameters were adjusted to 1 cycle of 45s at 98°C,14 cycles of 15s at 98°C followed by 10s at 60°C and 1 cycle of 1 min at 72°C. DNA was again purified using 1.1X AMPure XP beads and eluted in 0.1X TE buffer. Libraries were sequenced on the Illumina NextSeq platform at 41 bp paired-end.

### ChIP-seq

ChIP was carried out as previously described^19^., In brief, cells were grown to confluence and cross-linked in DMEM containing 1% formaldehyde for 10 min at room temperature, reaction was quenched with 200mM (final concentration) Glycine, cells were scraped off and rinsed with 10 ml 1 × PBS. Pellets were resuspended first in 10 ml buffer 1 (10 mM Tris (pH 8.0), 10 mM EDTA, 0.5 mM EGTA, 0.25% Triton X-100) and then in 10 ml buffer 2 (10 mM Tris (pH 8.0), 1 mM EDTA, 0.5 mM EGTA, 200 mM NaCl). Then cells were lysed in 1 ml lysis buffer (50 mM HEPES/KOH (pH 7.5), 500 mM NaCl, 1 mM EDTA, 1% Triton X-100, 0.1% DOC, 0.1% SDS, protease inhibitors) and sonicated for 20 cycles of 30s using a Diagenode Bioruptor Pico, with 30s breaks in between cycles. For the immunoprecipitation, lysate was first pre-cleared with protein A/G magnetic Dynabeads Magnetic beads (Thermo Fisher Scientific) for 1h at 4°C, then incubated with antibody of choice overnight at 4°C. The mixture was then incubated for 3h at 4°C with washed protein A/G magnetic Dynabeads Magnetic beads. Beads were washed three times with 1 ml lysis buffer, once with 1 ml DOC buffer (10 mM Tris (pH 8.0), 0.25 M LiCl, 0.5% NP-40, 0.5% deoxycholate, 1 mM EDTA), once with TE and bound chromatin was eluted in 1% SDS/0.1 M NaHCO_3_. After RNase A treatment, proteinase K digestion was performed at 55°C for 2h, before reversing the cross-linking by overnight incubation at 65 °C. DNA was isolated by purification using AMPure XP beads. A sample of the input chromatin was treated in the same way to generate total input DNA. Immunoprecipitated DNA and 200ng of input DNA were submitted to library preparation (NEBNext Ultra DNA Library Prep Kit, Illumina). In the library preparation protocol, input samples were amplified using 5 PCR cycles and immunoprecipitation samples using 12 cycles. Libraries were sequenced on the Illumina HiSeq 2500. All ChIP-seq experiments were performed in at least two independent biological replicates per condition.

### Co-IP MS protein enrichment analysis

Co-IP MS enrichment analysis was carried out as previously described^38^. In brief, protein identification and relative quantification was performed with MaxQuant (version 1.5.3.8) using Andromeda as the search engine^41^ and label-free quantification (LFQ)^42, 43^. The mouse subset of the UniProt version 2019_04 combined with the contaminant database from MaxQuant was searched and the protein and peptide FDR were set to 1% and 0.1% respectively. Protein intensities were first normalized to the smallest total sum of intensities across all samples, then log_2_ transformed after dividing samples by 2^20 and adding a pseudocount of 5 in order to stabilize variance of the data. Snf2h enriched samples were compared to datasets generated by co-IP MS in the Snf2hΔ line (i.e., mock IP) and significance estimates were determined using limma ^44^. Proteins with < 0.01 adjusted *P* value were considered significantly enriched.

### RNA-seq data analysis

RNA-seq reads were aligned to the mouse genome (mm10). Promoters were defined as ±1,000 nt around the TSS of each transcript in the UCSC Known Genes database, which was accessed via the Bioconductor package TxDb.Mmusculus.UCSC.mm10.knownGene (v 3.10.0). Reads were aligned using the qAlign function from the QuasR package (v 1.36.0), with parameters “splicedAlignment = TRUE” and “aligner = “Rhisat2”. Differential expression analysis was performed using gene-level quantifications and the quasi-likelihood method (glmQLFit and glmQLFTest functions) with default parameters using the edgeR package (v 3.40.2). First, weakly or non-detected genes were filtered out using the filterByExpr function, and then a model was fitted of the form ∼batch + genotype (where batch is a factor with levels corresponding to the batch of RNA-seq experiment associated with the sample, and genotype is a factor with levels corresponding to the genotype of the cell line). Differentially expressed genes (DEGs) for each KO-WT pair were selected among those having at least 2-fold change (absolute log_2_FC > 1) in either direction, and a FDR smaller than 0.01, as calculated by edgeR.

The same DEGs were clustered using kmeans (kmeans function from the stats package) with k = 4. The number of clusters, k, was determined by performing kmeans clustering over a range of k of 2-30 and selecting a value over which reduction in the total within-cluster sum of squares (wss) appeared less significant (“elbow method”). DEGs were displayed using ComplexHeatmap (v 2.12.0). For all plots the mean signal from at least two independent biological replicates is reported unless otherwise specified in the figure legend.

For Gene Ontology analysis, enriched “Biological Process” terms in both upregulated and downregulated genes for each genotype were searched using the enrichGO function in the clusterProfiler package (v 4.8.1), using all expressed genes as background and with a p-value cut-off of 0.01.

### ATAC-seq data analysis

ATAC-seq reads were trimmed using cutadapt ver2.5 with parameters -a CTGTCTCTTATACACA -A CTGTCTCTTATACACA -m 5 -overlap = 1, and mapped to the mm10 mouse genome using the qAlign function in QuasR with default parameters, which uses bowtie for short read alignments. ATAC-seq peaks were called using MACS2 (v 2.2.7.1) with parameters --nomodel --shift -100 --extsize 200 --keep-dup all -g mm --qvalue=1e-2. For comparative analysis, a unique peak set was created with all genomic regions that were called as a peak in at least two technical replicates of at least one sample.

Differentially accessible regions were called using read counts on peaks and the quasi-likelihood method (glmQLFit and glmQLFTest functions) with default parameters using the edgeR package (v 3.40.2). A model was fitted of the form ∼batch + genotype (where batch is a factor with levels corresponding to the batch of ATAC-seq experiment associated with the sample, and genotype is a factor with levels corresponding to the genotype of the cell line). Differentially accessible peaks were clustered using kmeans with k = 4 (using the kmeans function from the stats package). Log_2_ fold-changes in ATAC signal at these differentially accessible peaks were displayed using ComplexHeatmap (v 2.12.0).

For motif analysis, enrichment for each of the vertebrate transcription factor motifs contained in the JASPAR2022 database^45^ was calculated using the calcBinnedMotifEnrR in the monaLisa package^46^. For visualization, motifs that had a log_2_ fold-enrichment of > 1.5 and log10 adjusted p-value < 0.001 in at least one bin were selected.

ATAC–seq metaprofiles around bound CTCF sites were generated using the qProfile function from the QuasR package to get read counts in a 2-kb window anchored by the CTCF-binding motif, normalized by sequencing depth. Profiles were then smoothed with a running mean of 21 bp and multiplied by 100. For all plots the mean signal from at least two independent biological replicates is reported unless otherwise specified in the figure legend.

Strong DNAseI hypersensitive sites (DHS) were defined as previously described^14^.

### MNase-seq Data Analysis

MNase-seq reads were trimmed using cutadapt ver2.5 with parameters -a AGATCGGAAGAGCACACGTCTGAACTCCAGTCA -A AGATCGGAAGAGCGTCGTGTAGGGAAAGAGTGT -m 5 --overlap=1, and mapped to the mm10 mouse genome using the qAlign function in QuasR with default parameters, which uses bowtie for short reads alignment. MNase-seq metaprofiles around sites of interest (TSSs, DHSs or TF binding sites) were generated using the qProfile function from the QuasR Bioconductor package (v 1.26.0) with the parameter shift = ‘halfInsert,’ to get MNase fragment midpoint counts in a 2-kb window anchored to the TF-binding motif. Raw counts were normalized by dividing through the median of each profile and multiplying by the median of all sample medians. Profiles were then smoothed with a running mean of 21 bp. Similarly, heatmaps of MNase-seq fragment midpoints were generated by normalizing the profiles to RPKM.

Nucleosome repeat lengths were calculated using a Phasogram-based approach described in^14, 47^ implemented using the calcPhasogram and estimateNRL functions in the swissknife R package (https://fmicompbio.r-universe.dev/swissknife) with default parameters.

For plots showing MNase signal as a function of fragment length, we counted reads in a 2kb window centered on bound CTCF motifs (see ChIP-seq analysis) and divided based on the length of the sequenced fragment. Data was standardized using the scale function in R. For all plots the mean signal from at least two independent biological replicates is reported unless otherwise specified in the figure legend.

### CUT&RUN Data Analysis

CUT&RUN reads were trimmed using cutadapt (v 2.5) with parameters -a AGATCGGAAGAGCACACGTCTGAACTCCAGTCA -A AGATCGGAAGAGCGTCGTGTAGGGAAAGAGTGT -m10 -overlap = 1, and mapped to the mm10 mouse genome using the qAlign function in QuasR with default parameters. To account for differences in library size, the number of total reads mapped for each sample was scaled down to the sample with the lowest number of mapped reads. Average metaplots and single locus heatmaps were generated using the qProfile function in QuasR with default parameters; profiles were calculated over 2kb windows centered on either CTCF bound motifs or DHS center (see below) and smoothed over 51 bps. For the average metaplots the signal was divided by the total number of genomic regions considered. For the single locus heatmaps CTCF regions were sorted by Snf2h signal in WT. For the boxplots (Extended data), reads were counted over 250 bp windows centered on the region of interest using the QuasR function qCount, whereby reads were shifted by half the fragment length and each fragment was counted once. Log_2_ read counts were calculated as log_2_(n + 8), in which n are the library-size normalized counts and 8 is the pseudo-count, used to decrease noise levels at low read counts in any comparison. Enrichment over controls (IgG) were calculated by subtracting the log_2_ reads counts of the control from the log_2_ read counts of the corresponding sample. All plots were generated using the mean values from two independent replicates.

DNAseI hypersensitive sites (DHSs) and CTCF bound sites were defined as described in the above sections.

### ChIP-seq Data Analysis

ChIP-seq reads were trimmed using cutadapt (v 2.5) with parameters -a AGATCGGAAGAGCACACGTCTGAACTCCAGTCA -m 5 --overlap=1, and mapped to the mm10 mouse genome using the qAlign function in QuasR with default parameters, which uses bowtie for short reads alignment. ChIP enrichment between immuno-precipitated and input samples were calculated using:

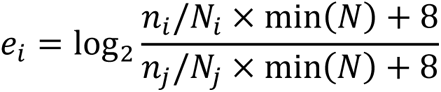

where *e_i_* is the ChIP enrichment of a region in sample *i*, *n_i_* and *n_i_* are the number of alignments in the immuno-precipitated sample *i* and the corresponding input sample *j*, *N_i_* and *N_j_* are the library sizes (total number of alignments) in samples *i* and *j*, and min(*N*) is the minimal library size over all samples. Changes of ChIP enrichment between two immuno-precipitated samples were calculated using the same formula. For genome-wide site predictions of CTCF, the motif MA0139.1 from the JASPAR2020 Bioconductor package was used. Bound CTCF sites were defined as motifs that have a log_2_(enrichment) (immunoprecipitation over input in a 251-bp window centered on the motif) of at least 1.0 (two-fold). CTCF motifs were clustered using kmeans with k = 5 (using the kmeans function of the stats package), using changes in ChIP-seq and ATAC-seq enrichment signal in BPTF ko cells compared to WT controls.

For the heatmap with average ChIP–seq profiles around per-cluster ATAC-seq peaks, ChIP–seq read counts in 2-kb windows centered on ATAC-seq peak midpoints were obtained using the qProfile function from the QuasR package pooling all samples measuring the same ChIP–seq target and normalizing them to RPKM. Normalized values were averaged across ATAC-seq peaks in each cluster and smoothed using a running mean of 45 bp. For better comparability between ChIP–seq targets, average cluster profiles from each target were further normalized by dividing through their maximum value or through 1.5 if the maximum was <1.5. For all plots the mean signal from at least two independent biological replicates is reported unless otherwise specified in the figure legend.

### Single-Molecule Footprinting analysis

Reads were trimmed using Trimmomatic v0.32^40^ in paired-end mode using the ILLUMINACLIP option. Trimmed reads were mapped to the mm10 mouse genome using the qAlign function from the QuasR package v_1.36.0 with parameters for bisulfite data. DNA methylation was quantified for all Cs using the qMeth function and then separated into Cs in the CpG or GpC context, removing GCG and CCG sequence contexts as these cannot be distinguished between endogenous methylation and SMF methylation. Plots of SMF data are of [1 − GpC methylation] to visualize the footprint.

### Deep learning model architecture, training and interpretation

A convolutional neural network (CNN) was trained on one-hot-encoded 150-bp long DNA sequence(s) centered at CTCF bound sites (n=47630) as input to predict the change in CTCF binding in BptfΔ cells compared to wild-type as measured by ChIP-seq. The architecture of CNN was adapted from Basset^48^ and further modified based on the DeepSTARR design by ref.^21^. The CNN in our study starts with four sequential convolutional layers (1D, filters = 128, 128, 128, 64; size = 5, 3, 5, 3) each followed by ReLU activation and max-pooling (size = 2). The output of the convolutional layers was fed into two fully connected layers with ReLU activation having 128 and 64 neurons respectively. Dropout of 0.4 was applied after each fully connected layer. The final layer was employed to predict the CTCF ChIP-seq changes in BptfΔ cells compared to wild-type, using a linear activation function. The model was implemented in the Keras framework^49^ using the Keras R package (v 2.2.5.0) with TensorFlow^50^ backend (v 2.0.0). The training was performed using a mean-squared-error loss function and the Adam optimizer^51^ with a batch size of 64, and monitored for early stopping based on validation loss (20% of training set) with patience of 15 epochs. CTCF sites from chromosomes 16, 17, 18 and 19 were excluded from the training (n=32988) and validation (n=8248) sets and kept as the test set (n=6394) for model evaluation. For model interpretability, the DeepExplainer implementation^52^ from the SHAP library^53, 54^ was used to calculate contribution scores for every nucleotide in the provided sequences around bound CTCF sites. As reference sequence for DeepExplainer, 100 dinucleotide-shuffled versions were generated for each CTCF site. To summarize the contribution of each nucleotide at each position across all input sequences, average contribution scores per position were computed for each of the four bases by taking the average of the contribution scores of the nucleotides present in the input sequence. The resulting contribution weight matrix (as introduced by ref^55^) was visualized using ggseqlogo^56^ (v 1.0). TFBSTools^57^ was used to identify the position of the M2 motif as defined by ref.^23^ (downloaded from CTCFBSDB 2.0^58^).

### Hi-C

Hi-C experiments were performed as previously described^14^. Hi-C reads were mapped allowing for split alignments to the mouse mm10 genome (BSgenome.Mmusculus.UCSC.mm10) using the Burrows-Wheeler Alignment Tool (bwa v 0.7.17) with parameters ‘bwa mem -SP5M’. Hi-C pairs were extracted from mapped files and processed using pairtools (10.5281/zenodo.5214125), specifically with the parse, sort and dedup functions with default parameters. Deduplicated pairs were converted to *cooler* format using cooler^59^ (v 0.8.7) using the ‘cload pairs’ function with default parameters. Matrices were then balanced keeping only cis-contacts using the cooler ‘balance’ function with default parameters. For all plots the mean signal from at least two independent biological replicates is reported unless otherwise specified in the figure legend.

### Public Datasets

ChIP–seq (metaprofiles and heatmaps at ATAC-seq peaks): H3K27me3: (GSE30203, samples GSM747539 to GSM747541) (ref.^60^), H3K4me1: (GSE30203, sample GSM747542) (ref.^60^), H3K27ac: (GSE67867, samples GSM1891651 and GSM1891652) (ref.^61^), H3K36me3: (GSE33252, samples GSM801982 and GSM801983) (ref.^62^).

## Acknowledgments

We thank members of the D.S. laboratory for their critical feedback on the study and manuscript. We thank Luca Giorgetti and Ilya Flamer for critical input on Hi-C experiments and analysis, Luke Isbel for help with CUT&RUN experiments, Ralph Grand and Marco Pregnolato for help with SMF experiments, Michael Stadler for advice on data analysis. We thank S. Smallwood and the functional genomics platform of the FMI for next-generation sequencing support, D. Hess and the proteomics and protein analysis platform of the FMI for the mass spectrometry support. D.S. acknowledges support from the Novartis Research Foundation, the Swiss National Science Foundation (310030B_176394 to D.S.) and the European Research Council under the European Union’s (EU) Horizon 2020 research and innovation programme grant agreements (ReadMe-667951 and DNAaccess-884664). M.Iurlaro was supported by a European Molecular Biology Organization advanced fellowship (ALTF 611-2019).

The funders had no role in study design, data collection and analysis, decision to publish or preparation of the manuscript.

## Authors contributions

M.Iurlaro, F.M. and D.S. conceived the study and planned the experiments. M.Iurlaro derived cell lines, performed the experiments and comprehensive data analysis. F.M. performed CUT&RUN and IP-MS experiments, and analyzed resulting data. C.W. performed co-IP and western blots experiments. M.Iskar performed deep learning modeling. L.B. supervised and instructed on data analysis. M.Iurlaro, F.M. and D.S. interpreted the results and wrote the manuscript with input from all authors.

## Extended Data Figures

**Extended Data Fig. 1.**
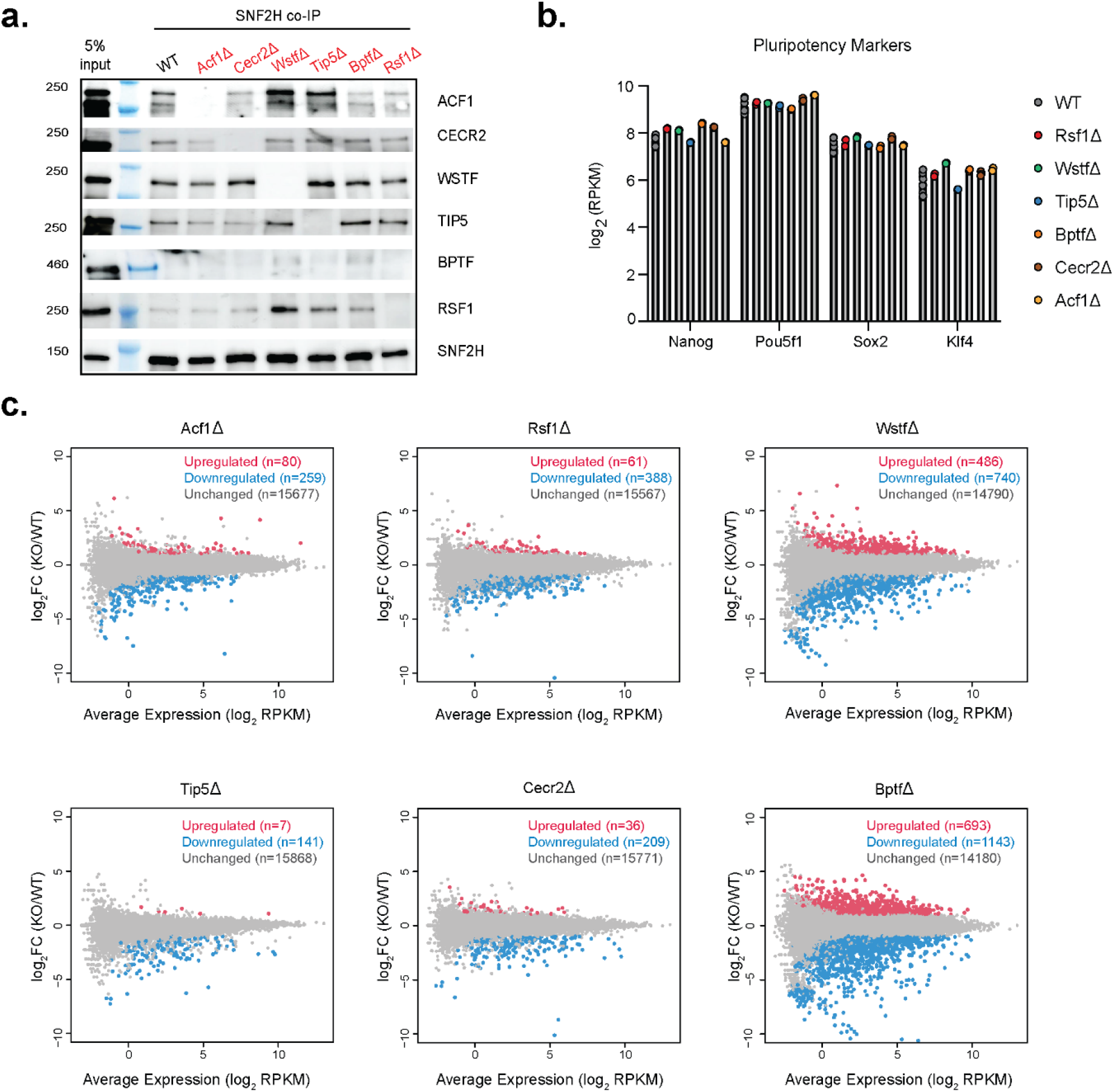
ISWI accessory subunit deletions are complex-specific and their absence does not affect pluripotency. **a.** SNF2H co-immunoprecipitation followed by western blot against ISWI subunits in WT and ISWI deletion lines (names highlighted in red for deletion lines). **b.** Expression of pluripotency genes (log_2_ RPKM) Nanog, Pou5f1(Oct4), Sox2 and Klf4 in WT and ISWI deletion lines. **c**. RNA changes in new deletion lines shown as MA plots. Differentially expressed genes are reported in red (upregulated) or blue (downregulated), unchanged genes are reported in gray.

**Extended Data Fig. 2.**
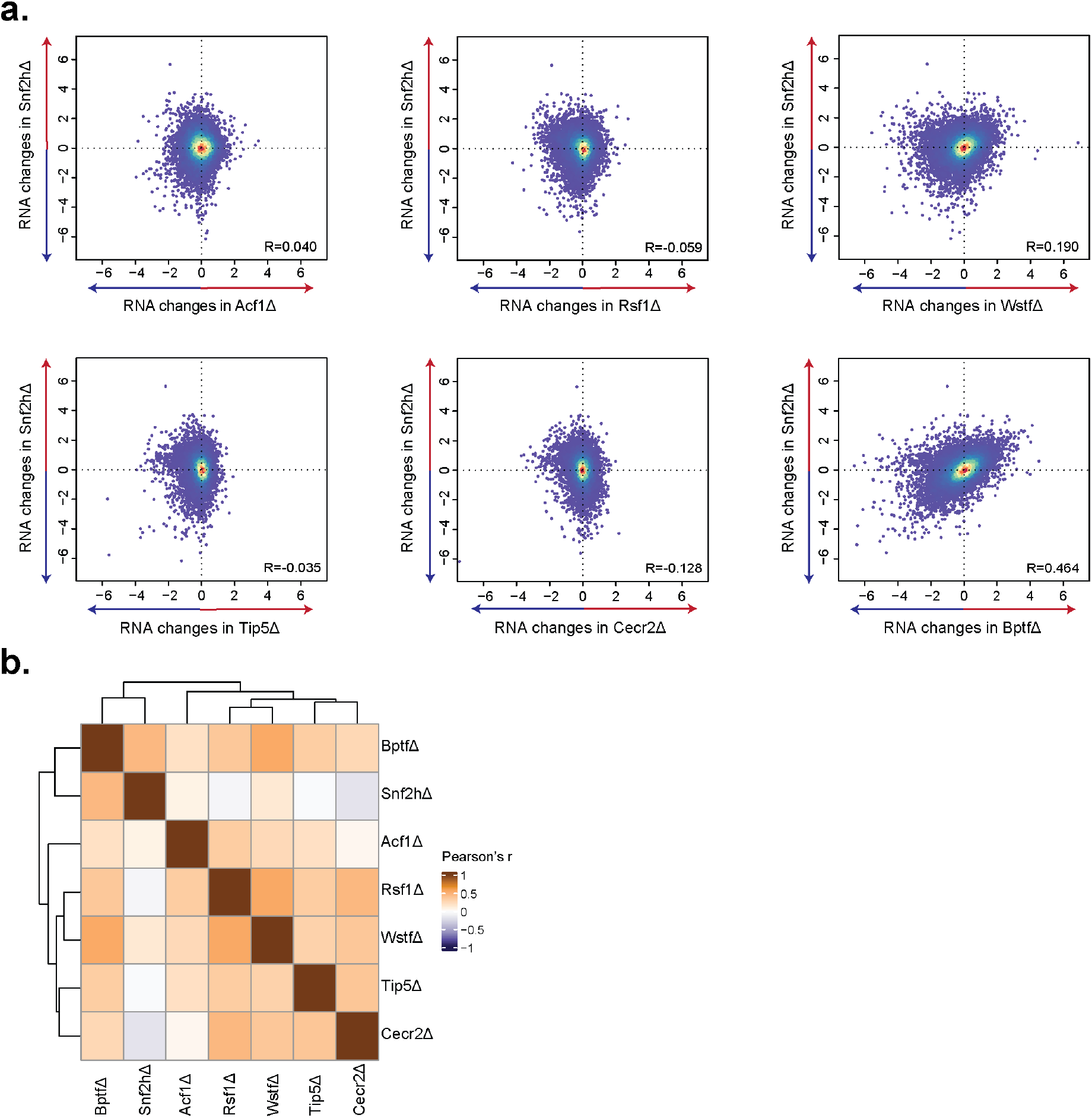
Bptf and Snf2h deletions present a similar transcriptional response. **a.** Quantitative comparison of RNA changes (log_2_FC) in accessory subunits deletions (x-axis) versus SNF2H deletion (y-axis). R: Pearson’s correlation coefficient. **b.** Heatmap of Pearson’s correlation of transcriptional changes induced by each deletion. Correlation was calculated on log_2_ fold-change data of genes called as differentially expressed in at least one contrast.

**Extended Data Fig. 3.**
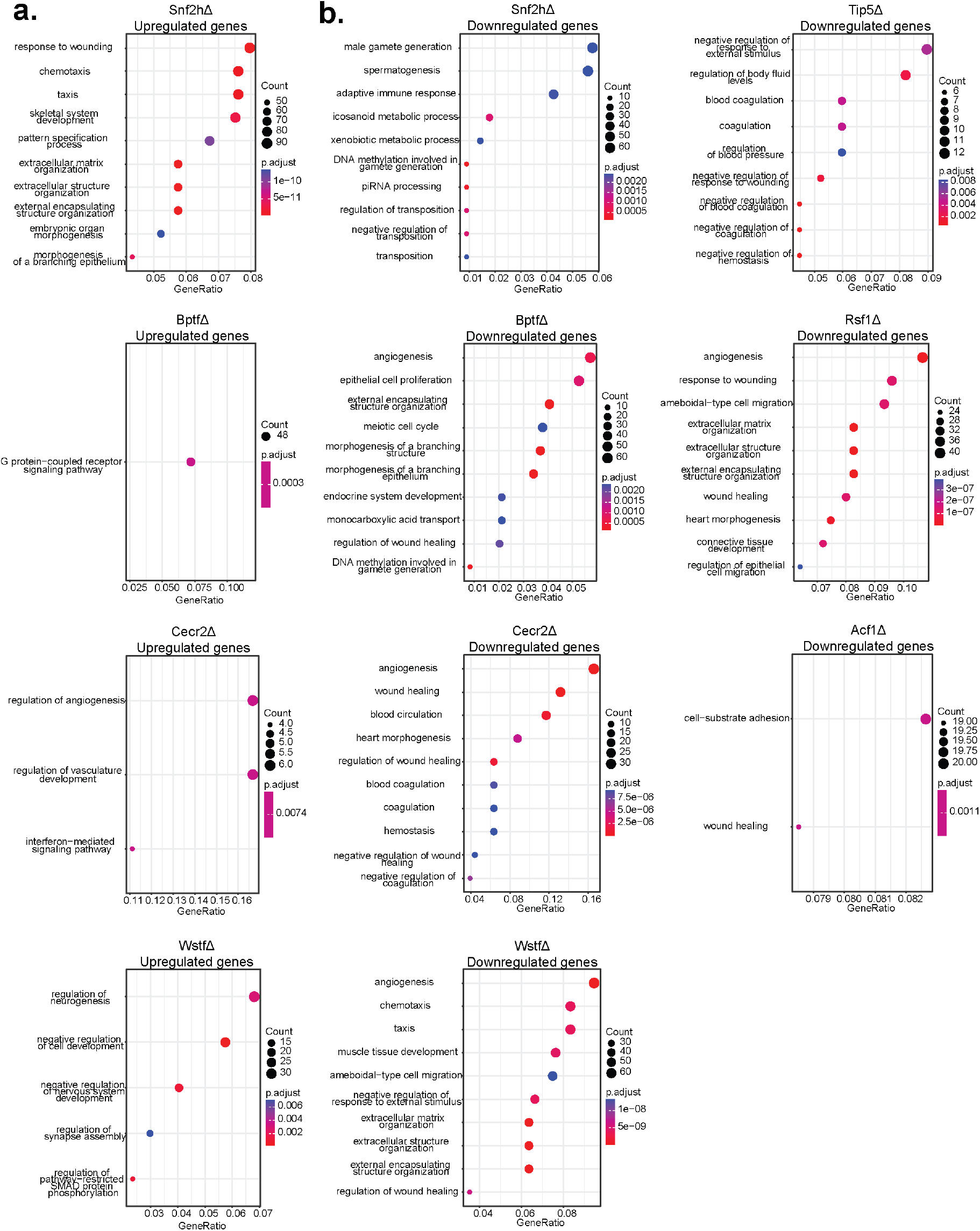
Gene Ontology analysis on differentially expressed genes in ISWI subunit deletions. **a.** Gene Ontology (see Methods) terms enriched in the set of upregulated genes of ISWI deletion lines. Adjusted p-value calculated through a one-sided Fisher’s exact test with Benjamini-Hochberg multiple testing correction. **b**. Gene Ontology terms enriched in the set of downregulated genes of ISWI deletion lines.

**Extended Data Fig. 4.**
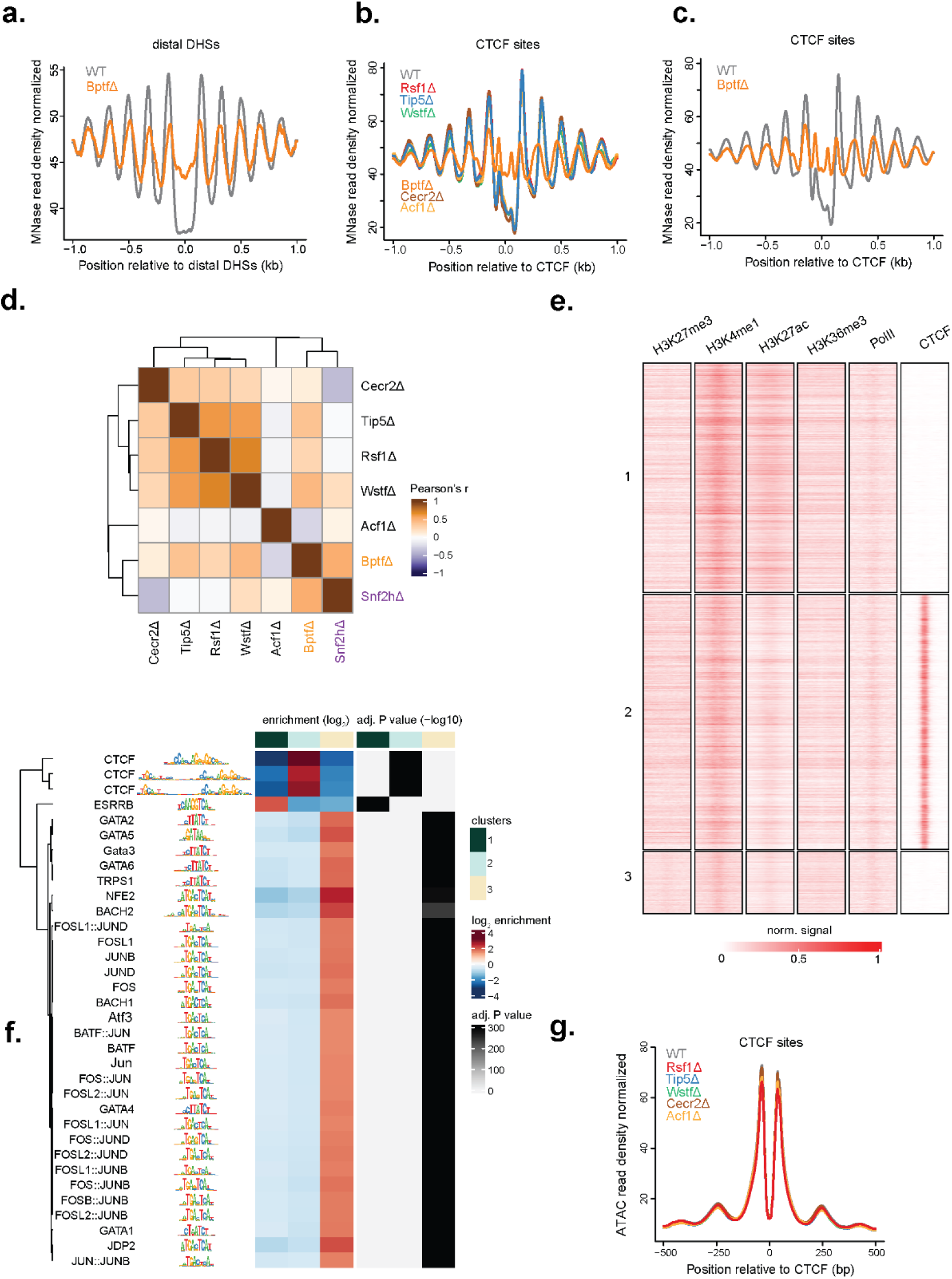
Nucleosome and accessibility profiling reveals a BPTF-specific response at CTCF sites. **a.** MNase average signal at distal DNaseI hypersensitive sites in WT and BptfΔ cells. **b.** Same analysis (as in **a**) for bound CTCF sites in WT and individual deletion lines. **c.** Same analysis (as in **b**) but displaying only WT and BptfΔ lines. **d.** Heatmap showing Pearson’s correlation of chromatin accessibility changes induced by each deletion. Correlation was calculated on log_2_ fold-change ATAC-seq signal on peaks called as differentially accessible in at least one comparison. **e.** Enrichment of chromatin marks and chromatin associated factors in clusters with differential accessibility response (clusters as in Fig. 2d are reported on the left). **f.** Motif enrichment analysis over the same clusters (as in **e**). Adjusted p-value calculated through a one-sided Fisher’s exact test with Benjamini-Hochberg multiple testing correction. **g.** Average ATAC-seq signal at bound CTCF sites in WT and individual deletion lines.

**Extended Data Fig. 5.**
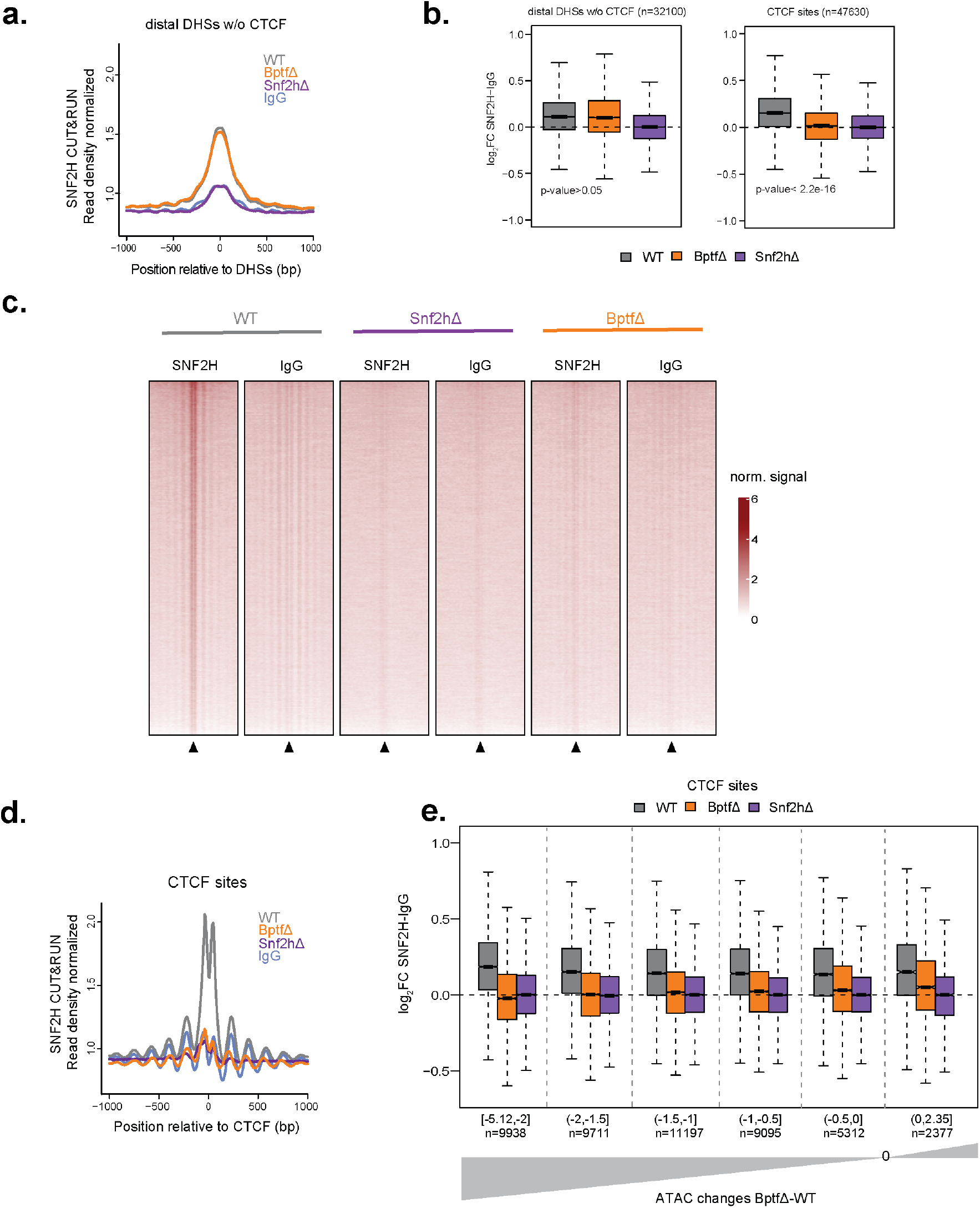
Absence of BPTF affects SNF2H localization at CTCF sites. **a.** Detection of SNF2H by CUT&RUN at distal DNaseI hypersensitive sites shown as average signal for WT (gray), BptfΔ (orange), Snf2hΔ (purple). IgG signal is shown in blue as negative control. CTCF sites are excluded to illustrate consistent binding of SNF2H in WT and BptfΔ lines. **b.** Boxplots showing SNF2H CUT&RUN signal (log2 fold-change over IgG control) for WT (gray), BptfΔ (orange), Snf2hΔ (purple) at distal DNaseI hypersensitive regions (left), and at bound CTCF sites (right) illustrating BPTF specific reduction in SNF2H signal over CTCF sites. Significance between WT and BptfΔ conditions was calculated by a one-sided Wilcoxon signed rank test. **c.** CUT&RUN signal for SNF2H in WT, BptfΔ, Snf2hΔ cells, shown as alignment densities centered on CTCF bound motifs (black arrow). IgG signal is shown for each background as negative control. **d.** SNF2H CUT&RUN average signal at bound CTCF sites for WT, BptfΔ, Snf2hΔ and IgG. **e.** Boxplot showing SNF2H CUT&RUN signal (log2 fold-change over IgG control) for WT (gray), BptfΔ (orange), Snf2hΔ (purple) at bound CTCF sites grouped by ATAC changes in BptfΔ. Groups are displayed from left to right starting with regions with stronger ATAC loss on the left (n= number of sites in each group).

**Extended Data Fig. 6.**
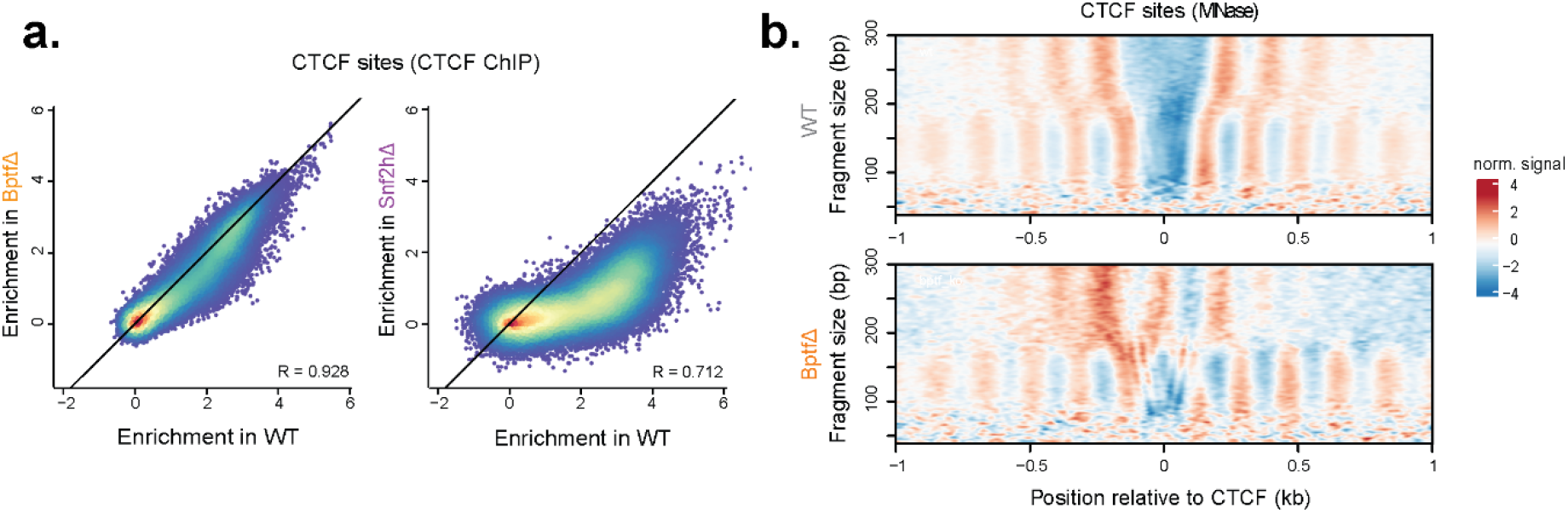
CTCF binding largely persists in absence of BPTF but coincides with changes in nucleosome organization. **a.** Quantitative comparison of CTCF binding (log_2_ fold-change IP/input) in WT cells (x-axis) vs BptfΔ (left) or Snf2hΔ (right) (y-axis), illustrating persistent binding in BptfΔ versus the drastic reduction in Snf2hΔ. R: Pearson’s correlation coefficient. **b.** V-plots representing scaled MNase data of fragment size on the y-axis and fragment midpoint position on the x-axis at bound CTCF sites in WT and BptfΔ cells, highlighting relative accumulation of longer MNase fragments (> 200bp) spanning CTCF bound sites upon deletion of BPTF.

**Extended Data Fig. 7.**
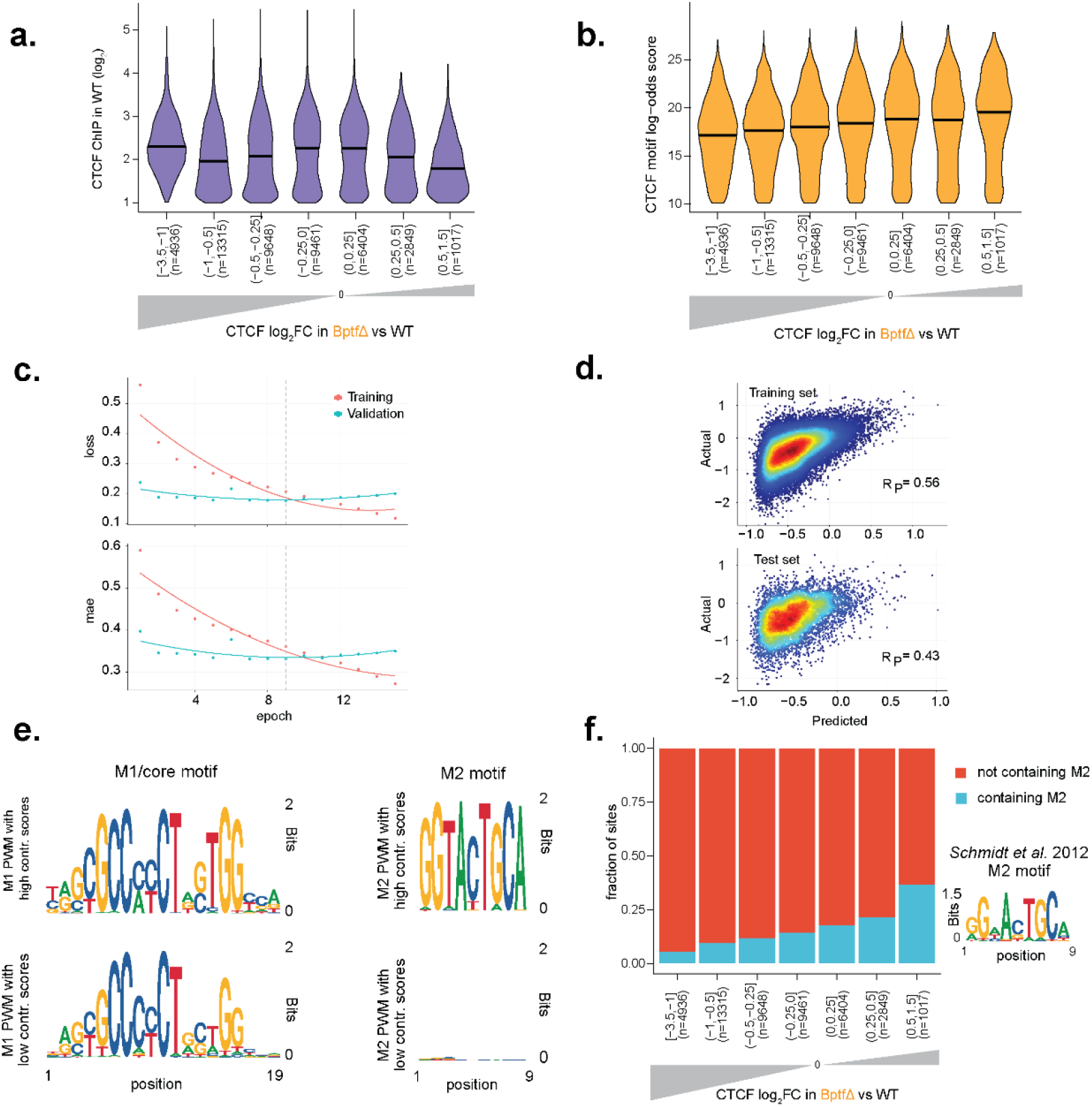
Deep learning identifies CTCF motif features enriched at sites of persistent binding in absence of NURF. **a.** CTCF sites were grouped based on changes in CTCF binding in BptfΔ versus WT. For each group the binding strength in WT cells is shown as violin plots with median (black line). n= number of CTCF sites within each group. **b.** Same groups (as in **a**) but now showing CTCF motif score (canonical motif M1 log-odds score) as violin plots with median (black line). Illustrating a trend for higher motif score in sites with persistent CTCF binding in absence of Bptf. **c.** Plots of loss (mean squared error, top) and the mean absolute error (bottom) metrics at each training epoch step for the training (red line) and validation sets (blue line). The dotted line indicates the selected epoch with the minimum validation loss. **d.** Scatter plot showing the observed vs predicted CTCF ChIP-seq log_2_ fold-change in BptfΔ compared to WT for the training (top) and the test set from held-out chromosomes (bottom). R_p_ indicates the Pearson correlation coefficient. **e.** Position weight matrix logos were generated in bits for the CTCF sites with the highest (n=1000, top) and lowest (n=1000, bottom) contribution scores calculated from the deep learning model. Sequence logos were created independently for M1 (left) and M2 (right) motifs. **f.** Fraction of CTCF sites containing an M2 motif (as defined by ref.^23^) and grouped (as in **a** and **b** by CTCF changes in BptfΔ), illustrating the increased presence of M2 at sites with persistent CTCF binding in absence of BPTF.

**Extended Data Fig. 8.**
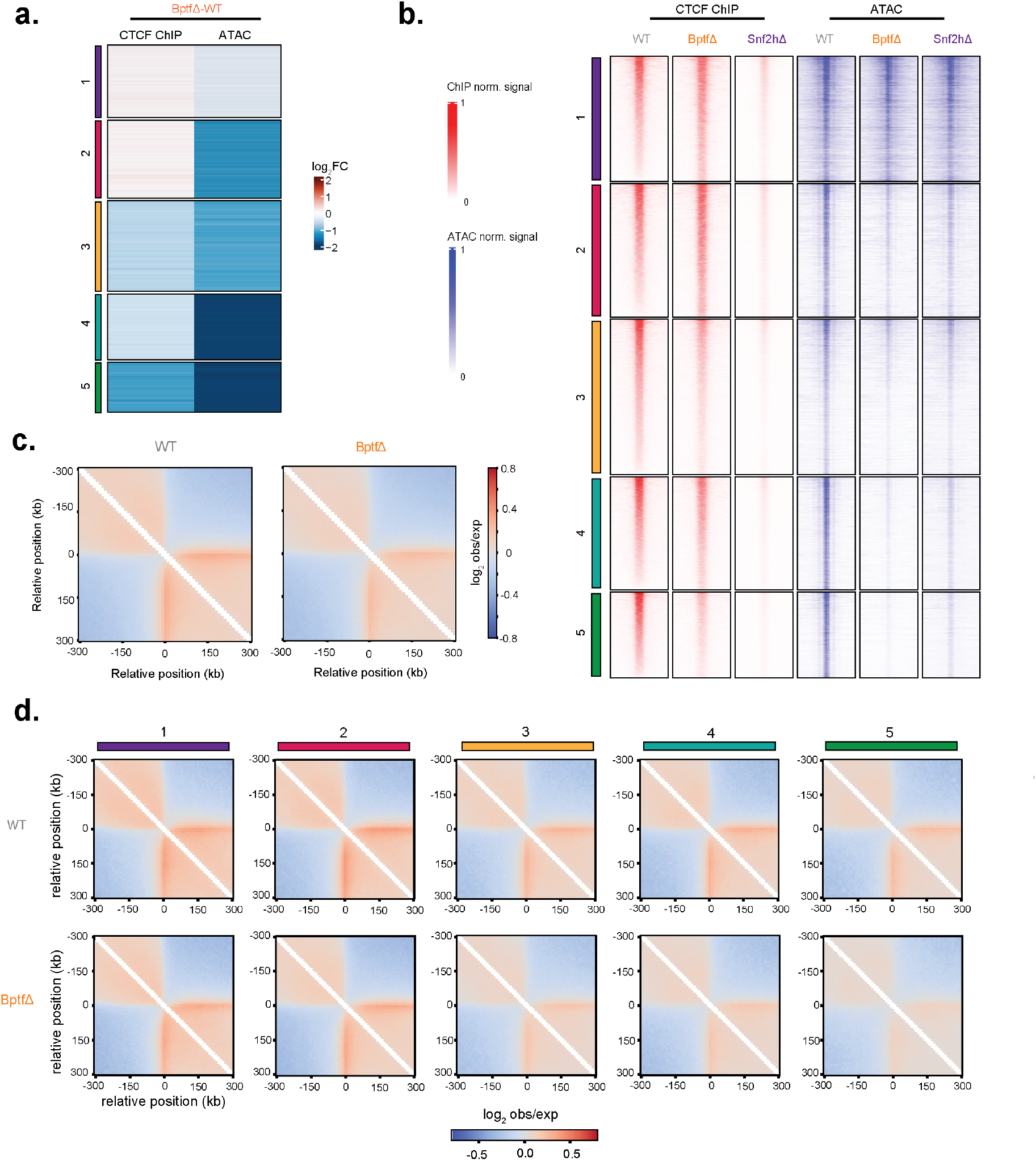
Absence of BPTF causes changes in long range chromatin contacts. **a.** Heatmap showing log_2_ fold-changes in CTCF binding (ChIP-seq) and accessibility (ATAC-seq) upon BPTF deletion at bound CTCF sites clustered (as in Fig. 4a). Cluster numbers reported on the left. **b.** CTCF binding (ChIP-seq) and chromatin accessibility (ATAC-seq) signal at clustered bound CTCF sites (as in **a**), in WT, BptfΔ, and Snf2hΔ cells. Cluster numbers reported on the left. **c.** Observed/expected interactions at bound CTCF sites in WT and BptfΔ cells, measured using Hi-C. **d.** Same (as in **c**), with CTCF sites divided into clusters (as in **a** and **b**). Cluster numbers reported on top.

**Extended Data Fig. 9.**
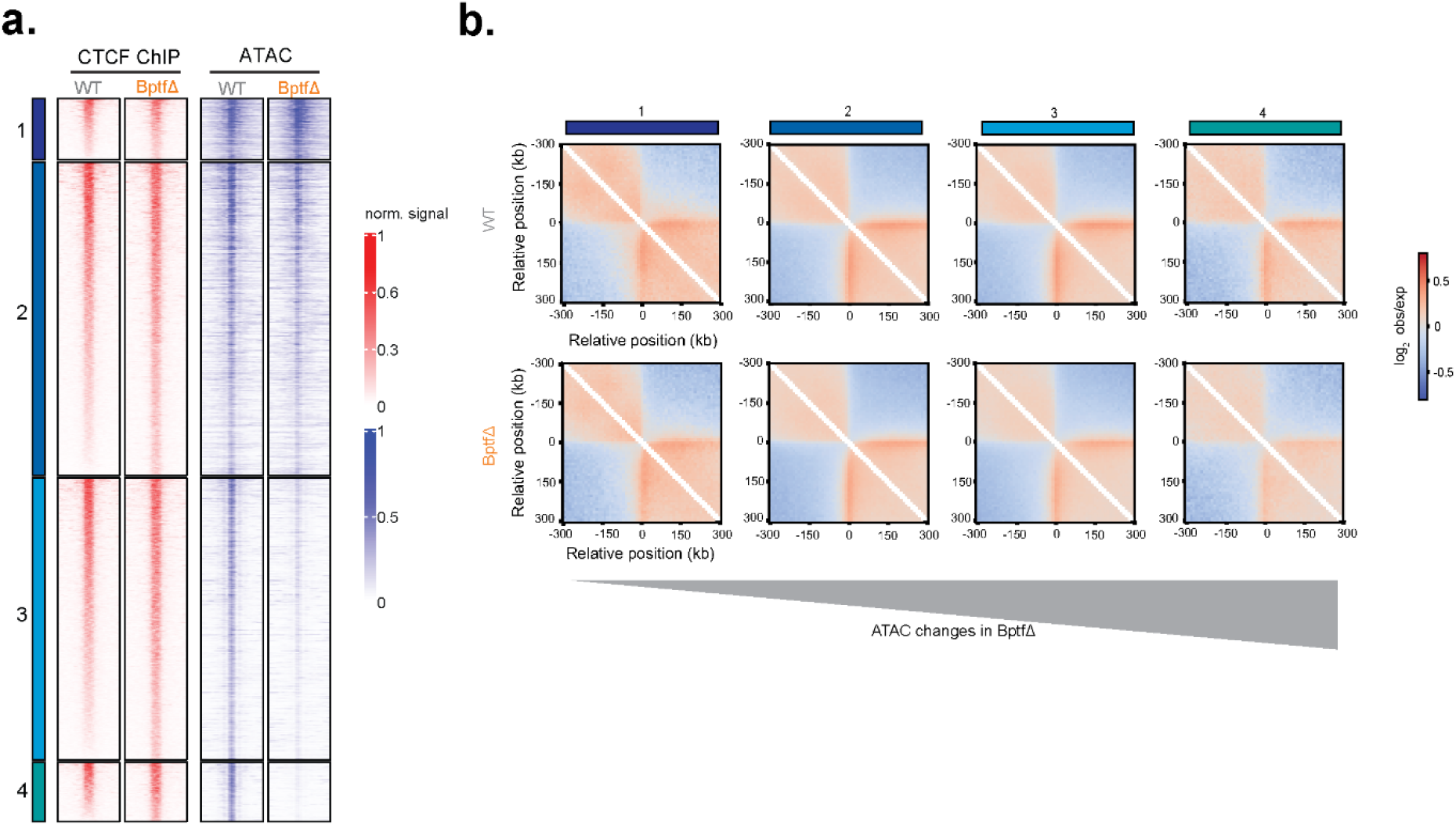
NURF dependent accessibility loss relates to loss of long range chromatin interactions. **a.** CTCF binding (ChIP-seq) and chromatin accessibility (ATAC-seq) signal in WT and BptfΔ cells within the four groups defined (as in Fig. 5a). Group numbers reported on the left. **b.** Observed/expected interactions within the same groups (as in **a**) measured using Hi-C in WT and BptfΔ cells. Group numbers reported on top.

